# A novel KIR3DL2-directed CAR T-cell therapy for cutaneous and systemic T-cell lymphomas

**DOI:** 10.64898/2026.11.05.764385

**Authors:** Federico Mario Aletti, Jinsheng Weng, Cao Cuong Le, Jingwei Liu, Faezeh Darbaniyan, Abdur Rehman, Matthew D. Richard, Selin Kolsuz, Xiaoyun Cheng, Sridevi Patchva, Yongfu Tang, Chun-Yen Lin, JingJing Cao, Kamini Singh, Shriya D. Reddy, Shiv G. Rawat, Swati Mohapatra, Jinsong Wei, Karan L. Chohan, Claudia Kettlun, Nicholas J. Dcunha, Neeraj Saini, Fuliang Chu, Michael R. Green, Swaminathan P. Iyer, Francisco Vega, Sattva S. Neelapu

## Abstract

Chimeric antigen receptor (CAR) T-cell therapy has transformed the treatment of B-cell malignancies and multiple myeloma, but its extension to T-cell lymphomas (TCLs) has been constrained by the paucity of antigens that can be safely targeted. While targeting pan-T antigens broadens the therapeutic reach across various TCLs, it carries a substantial risk of immunosuppression and infection. Here, we report killer cell immunoglobulin-like receptor, three Ig domains and long cytoplasmic tail 2 (KIR3DL2/CD158k) as a viable target for CAR T-cell therapy against a wide range of TCLs. Using single-cell RNA-seq data analysis and flow cytometric phenotyping, we observed that KIR3DL2 expression is restricted to a subset of mature NK cells and a minor fraction of CD8⁺ T cells, with negligible expression in other healthy adult tissues. Among TCLs, KIR3DL2 expression was noted in various subtypes of cutaneous and systemic TCLs. Anti-KIR3DL2 CAR constructs (3DL2CARs) efficiently and selectively redirected T cells against antigen-positive cell lines but not isogenic KIR3DL2-negative counterparts, with antigen-specific signaling, cytotoxicity, proliferation, and cytokine production. Importantly, 3DL2CART cells produced sustained tumor regression and prolonged survival in two distinct cell line-derived xenograft models of cutaneous TCL and in one patient-derived xenograft model of hepatosplenic TCL. 3DL2CART cells exhibited minimal off-target activity against bystander immune cells and showed no appreciable fratricide. To mitigate the risk of tumor cell contamination in autologous CAR T-cell products, we established a two-tiered strategy that integrates immunomagnetic depletion of KIR3DL2-positive tumor cells with *in vivo* activation of a safety switch if inadvertently transduced malignant cells contribute to immune escape. Together, our results establish the preclinical efficacy and safety of 3DL2CART-cell therapy and support its clinical translation. This approach may offer a targeted alternative to pan-T-cell-directed CAR T-cell strategies for patients with relapsed/refractory cutaneous and systemic TCLs, addressing a critical unmet clinical need.

## INTRODUCTION

T-cell malignancies account for 10–15% of lymphoid neoplasms^1^ and generally carry poorer prognoses and fewer treatment options than B-cell neoplasms^2–5^. Chimeric antigen receptor (CAR) T-cell therapy has transformed standards of care and outcomes in B-cell malignancies and multiple myeloma (MM) in recent years^6–15^ but is still far from delivering comparable results in T-cell lymphomas (TCLs). The challenge for development of CAR T-cell therapy in T-cell malignancies has always been the identification of antigens^16^. So far, most approaches have focused on pan-T-cell antigens (e.g., CD7, CD5), which are broadly shared between effector and target cells^17–20^. Potential risks with this approach include induction of widespread T-cell aplasia and sustained immunodeficiency in the host^21,22^; consequently, most pan-T-targeting products can only be used as a bridge to allogeneic transplantation, adding cost, toxicity, and risk for the patient^23,24^. Moreover, *trans* and *cis* interactions between the binder and its cognate antigen within and between CAR T cells may cause fratricide^17,19^, impair CAR surface expression^25^, and lead to early exhaustion or activation-induced cell death (AICD) during manufacturing^26^. These can be overcome by antigen downregulation on the therapeutic product, at the price of increased lentiviral cargo size^27^ or manufacturing complexity and/or potential genotoxicity if a nuclease-based knockout approach is used^28^. Lastly, in the setting of leukemic phase, the difficulty of separating malignant from healthy cells within the apheresis product^29^ creates a risk of inadvertent CAR transduction of tumor cells, which can result in *cis* antigen masking and subsequent immune escape^30^.

Killer cell immunoglobulin-like receptor, three Ig domains and long cytoplasmic tail, 2 (KIR3DL2/CD158k) is an inhibitory killer-cell immunoglobulin-like receptor involved in NK cell licensing. It binds to the HLA-A3 and HLA-A11 allotypes^31^, the β2-microglobulin-free heavy-chain homodimer of HLA-B27^32^, and CpG oligodeoxynucleotides (which it shuttles to endosomal TLR9^33^). KIR3DL2 expression is largely restricted to a subset of NK cells and rare T cells in healthy individuals^34,35^, whereas it is aberrantly expressed in some aggressive cutaneous TCL (CTCL) subtypes, such as Sézary syndrome (SS) and transformed mycosis fungoides (MF)^36,37^, as well as some cases of primary cutaneous anaplastic large cell lymphoma (pcALCL)^38^. Beyond CTCL, KIR3DL2 overexpression has been reported in up to 38% of systemic TCLs, including hepatosplenic TCL (HSTL), monomorphic epitheliotropic intestinal TCL (MEITL), extranodal NK/TCL (ENKTL), large granular lymphocytic leukemia (LGL), adult T-cell leukemia/lymphoma (ATLL), peripheral TCL-not otherwise specified, and in a minority of other subtypes^39,40^. TCLs are rare, mostly aggressive malignancies with limited durability of responses beyond first-line therapy and poor overall survival^41,42^. An effective CAR T-cell therapy targeting KIR3DL2 would address a substantial unmet need in relapsed/refractory (r/r) cutaneous and systemic TCLs, while preserving the advantages of targeting an antigen with largely tumor-restricted expression.

## METHODS

### Cell lines

HUT-78 (SS), HH (primary cutaneous TCL), MAC-2A (pcALCL), CA46 (Burkitt lymphoma), and HEK293T cell lines were purchased from the American Type Culture Collection (ATCC). Jurkat-Lucia™ NFAT reporter cell line was purchased from InvivoGen. All cell lines were cultured following the supplier’s recommended protocols. All lymphoma target/non-target cell lines were lentivirally transduced to express firefly luciferase (ffluc) and enhanced green fluorescent protein (eGFP) or mCherry. Isogenic KIR3DL2^+^ HH (HH^KIR3DL^^2^) were generated by lentiviral delivery of the full-length human KIR3DL2 protein. Isogenic, KIR3LD2^-^ subclones of HUT-78 were generated using CRISPR/Cas9 knock-out (KO). Single-guide RNAs and recombinant SP Cas9 V3 were purchased from Integrated DNA technologies (IDT) and delivered to the cells though Neon™ electroporation (InvitroGen) following the manufacturer’s protocol.

### CAR constructs generation

A proprietary murine anti-KIR3DL2 monoclonal antibody (mAb, clone K36) was generated through hybridoma technology (Suppl. methods). Clone K36 variable light (VL) and variable heavy (VH) chains were connected by a flexible Gly-Ser linker and single-chain variable fragment (scFv)-derived 3DL2CARs were cloned into a lentiviral backbone under the control of the murine stem cell virus (MSCV) promoter. 3DL2CARs included CD8α, CD28 or IgG4 hinge (H) fused with CH3 domain and CD8α or CD28 transmembrane (TM) domain, CD28 or 4-1BB co-stimulatory (CO) domains, and CD3ζ signaling domain, a 2A ribosomal skip element, and a truncated B-cell maturation antigen lacking the cytoplasmic signaling domain but with two copies of the extracellular domain fused in tandem (chimeric dual truncated BCMA or cdtBCMA) as a transduction marker/safety switch^43^. A non-signaling control CAR was obtained by truncating the CAR construct after the transmembrane domain.

### Lentivirus generation

Lentivirus (LV) for CAR transduction was generated by transfection of 80-100% confluent HEK293T cells with the transfer, packaging, and envelope plasmids (ABM 3rd Generation Packing System Mix) using Lipofectamine 3000 transfection kit (Invitrogen). LV supernatant was harvested at 24 and 48 hours post transfection, centrifuged and concentrated (Lenti-X-Concentrator, Takara Bio) at 1500xg for 45min at 4°C, re-suspended in Opti-MEM medium (Gibco) and used either fresh or cryopreserved at -80°C.

### CAR T-cell generation

Buffy coats from healthy donors were sourced from the local blood bank. Peripheral blood mononuclear cells (PBMCs) were isolated using density gradient separation and cryopreserved in aliquots with CryoStor CS10 freezing medium (STEMCELL Technologies). Thawed PBMCs were cultured overnight in complete medium supplemented with IL-2 (300 IU/mL, Cell Signaling). Primary T cells were isolated from PBMCs using the Pan T-cell isolation kit (Miltenyi Biotec), resuspended in complete medium at a concentration of 1×10⁶ cells/mL and activated using ImmunoCult Human CD3/CD28/CD2 T-Cell Activator (STEMCELL Technologies). After 48h, activated T cells were re-suspended in Opti-MEM medium supplemented with IL-2, plated in 24-well plates at 1×10⁶ cells/well in the presence of 10 µg/mL Vectofusin-1 (Miltenyi Biotec) and spin-transduced with lentivirus at 800xg for 1.5 hours at 32°C. Transduced T cells were expanded in complete medium supplemented with 300 IU/mL of human IL-2 and split every 1-2 days to maintain a concentration of ≤1×10⁶ cells/mL. Transduction efficiency was evaluated by flow cytometry on days 3 and 7 post-transduction. CAR T cells were used for functional assays either fresh or cryopreserved on day 7 post-transduction. Untransduced (UTD) donor-matched T cells were similarly activated and expanded *in vitro*.

### Flow cytometry

Cells were resuspended in staining buffer (1X phosphate-buffered saline [PBS] with 2% fetal bovine serum [FBS]) and pre-incubated with FcR Blocking Reagent (Miltenyi Biotec) for 10 min at room temperature. Cells were then incubated at 4°C for 30 min with the appropriate fluorescently labeled antibodies (BioLegend, BD Biosciences, ACROBiosystems). Antibodies used for flow cytometric analysis include CD2, CD3, CD4, CD8, CD19, CD20, CD14, CD16, CD56, BCMA, CD45, CD45RA, CD62L, HLA-A2, HLA-A3, FMC63, His tag, and IgG2a kappa isotype control. KIR3DL2 expression was detected with AF647-conjugated clone K36 (Suppl. Methods). Expression of the anti-KIR3DL2 scFv was detected using histidine-tagged recombinant human KIR3DL2 protein (TargetMol), followed by a fluorescently labeled anti-His tag antibody. Cell viability was assessed using LIVE/DEAD Fixable Dead Cell Stain Kits (Thermo Fisher Scientific). Antigen density on target cell lines was assessed by Quantum Simply Cellular (QSC) microspheres (Bang Laboratories). Samples were acquired on a BD LSRFortessa flow cytometer (BD Biosciences) or a Cytek Aurora spectral cytometer (Cytek Biosciences) and analyzed using FlowJo v10 software (Tree Star).

### Cytotoxicity, proliferation and cytokine release assays

For cytotoxicity assays, eGFP or mCherry-transduced target cells were co-cultured with effector cells at the specified effector-to-target (E:T) ratios using the Incucyte Live-Cell Analysis System (Sartorius). The fold change in live tumor cells normalized to baseline (Tumor Index) was monitored every 6 hours by measuring the mean green/orange area confluence. For proliferation assays, CAR T cells were labeled with CellTrace™ Far Red (Thermo Fisher Scientific) and co-cultured with eGFP-positive target cells at 1:1 E:T ratio for 96h. Proliferation was assessed by flow cytometry by estimation of mean fluorescence intensity change (MFI) of CellTrace™ Far Red-labeled T-cell population. Cell culture supernatants from the cytotoxicity assays were collected 48 hours after co-culture and analyzed for cytokine levels using the Meso Scale Discovery multiplex platform (Meso Scale Diagnostics). In all functional assays, the number of effector T cells was normalized to the lowest CAR expression level by adding UTD T cells where appropriate, so that the number of CAR-positive and total T cells per well was kept constant.

### In vivo studies

All animal experiments were conducted in compliance with protocols approved by the Institutional Animal Care and Use Committee (IACUC) of the University of Texas MD Anderson Cancer Center. Ffluc tumor cells were intravenously injected into 6/8-week-old NSG mice (NOD.Cg-Prkdc^^scid^Il2rg^^tm1^Wjl/SzJl, Jackson Laboratories). Cryopreserved CAR T cells were normalized for CAR expression and total T cell number by adding donor-matched UTD T cells and administered as a single intravenous injection after confirming tumor engraftment. Tumor burden was assessed using the Xenogen IVIS Lumina imaging system (Caliper Life Science) after intraperitoneal injection of 200 uL of 15mg/mL D-Luciferin (PerkinElmer). Luminescence data were normalized and analyzed using Living Image software (PerkinElmer). CBTL-81777 (HSTL patient-derived xenograft [PDX]) was purchased from The Public Repository of Xenograft (PRoXe), injected via tail vein and expanded in NSG mice. Upon reaching humane endpoints, spleen and liver were harvested, processed into single-cell suspensions, and cryopreserved in CryoStor CS10 freezing medium (STEMCELL Technologies) for long-term storage.

### Statistical analysis

Single-cell RNA sequencing (scRNA-seq) data describing KIR3DL2 expression across healthy human tissues and cell types were obtained from The Human Protein Atlas https://www.proteinatlas.org/ version 22) and analyzed in R (version 4.6.1) using RStudio (version 2026.09.0, Posit). Other statistical analyses were performed using GraphPad Prism version 8.0. Significance for functional assays was determined by Student’s t test, one-way or two-way ANOVA followed by appropriate post-hoc tests, as indicated in the figure legends. Survival analyses were performed using the Kaplan-Meier method, and differences between groups were evaluated by log-rank tests with Holm-Bonferroni adjustment for multiple comparisons.

## RESULTS

### KIR3DL2 expression pattern in healthy tissues and immune subsets is favorable for CAR T-cell targeting of different TCL subtypes

Analysis of publicly available scRNA-seq data from The Human Protein Atlas confirmed transcript expression well above the detection threshold restricted primarily to NK cells and, to a lesser extent, T cells. Negligible or no expression was found on other immune subsets and cell types (**Fig. 1A**). We phenotyped PBMCs from 30 healthy donors and confirmed expression at protein level restricted to ∼10–50% of NK cells, largely confined to the mature CD56^dim^CD16^+^ subpopulation, and to 10-50% of CD8^+^ terminally differentiated effector memory (TEMRA) cells, with marked variability among donors. The remaining CD8^+^ and CD4^+^ T cells, B cells and monocytes exhibited low or undetectable levels (**Fig. 1B**). Elevated KIR3DL2 expression was confirmed by immunohistochemistry (IHC) in 32/50 (64%) of TCL biospecimens tested, including 7/8 (88%) CTCL cases, as well as HSTL, ENKTL, nodal γδTCL (nγδTCL), primary cutaneous γδTCL (pcγδTCL), MEITL, ATLL and others (**Fig. 1C, D** and **Suppl. Fig. 1A**; Suppl. Methods). Flow cytometric analysis confirmed endogenous KIR3DL2 expression in HUT-78 (SS cell line) with low antigen density (∼3,500 molecules/cell) and MAC-2A (pcALCL, ∼4,500 molecules/cell), as well as in HH^KIR3DL^^2^ (CTCL cell line, endogenously negative and with ∼20,000 molecules/cell upon lentiviral delivery) (**Fig. 1E** and **Suppl. Fig 1B**).

**Figure 1.**
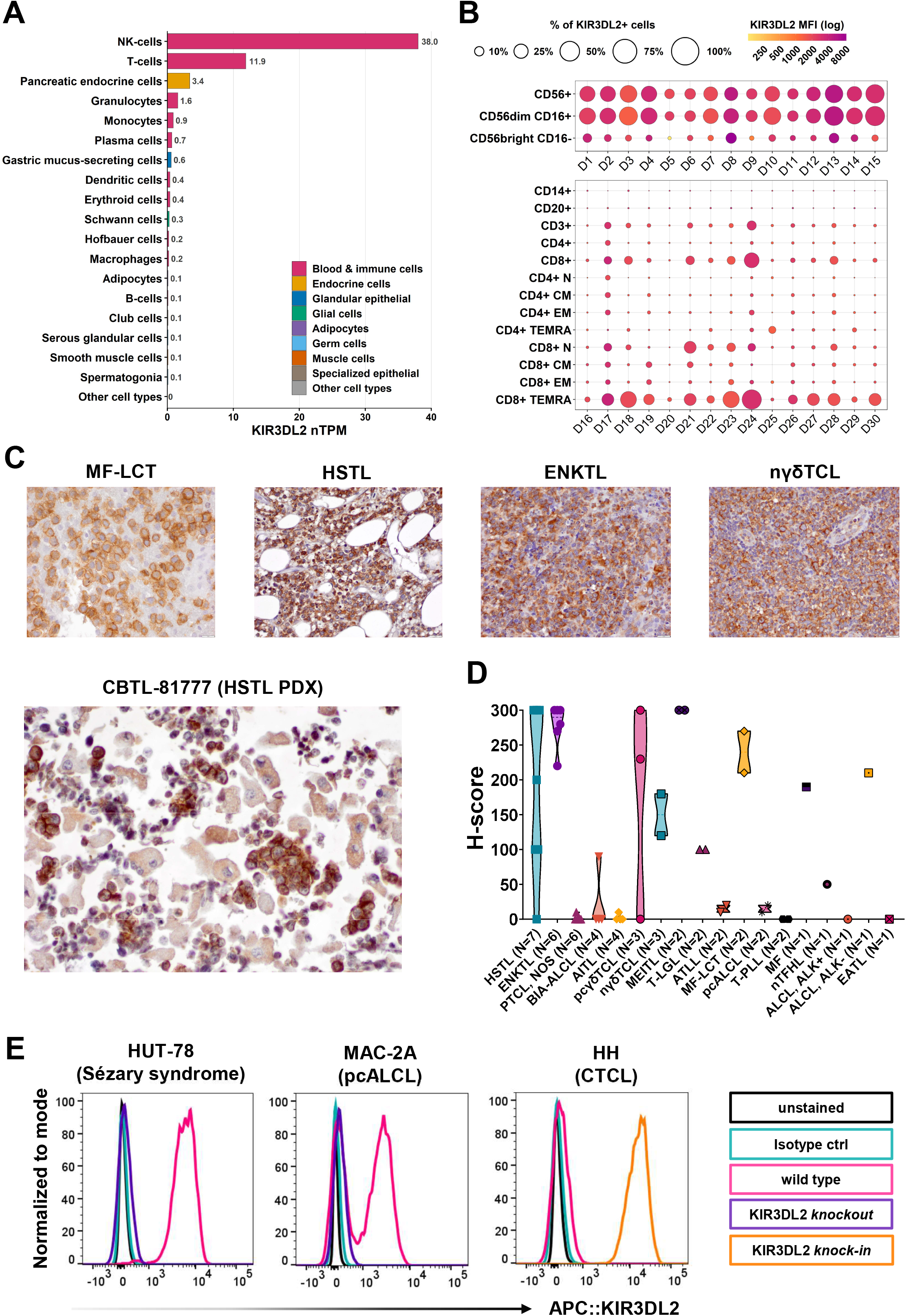
KIR3DL2 expression pattern in healthy tissues and immune subsets is favorable for CAR T-cell targeting of a wide range of TCL subtypes. **A)** Single-cell RNA sequencing (scRNA-seq) data from 80 cell types across 29 human tissues and peripheral blood mononuclear cells (PBMCs) were screened for KIR3DL2 expression. All cell types with nTPM > 0 are reported in descending order. Source: The Human Protein Atlas v22.0. **B)** Bubble plot representing KIR3DL2 protein expression detected by flow cytometry across different immune subsets from 30 healthy donor PBMCs. Top panel: NK cell subsets (n = 15 donors). Bottom panel: monocytes, B cells and T cells, including memory subsets, in a separate cohort of healthy donors (n = 15). Dot size represents the percentage of cells expressing KIR3DL2 and color intensity represents the log-transformed MFI within each subset. **C)** Representative immunohistochemistry (IHC) pictures for KIR3DL2 in patient biospecimens of MF-LCT, HSTL, ENKTL, nγδTCL and a HSTL patient-derived xenograft (KIR3DL2-positive). **D)** Violin plot of H-scores for KIR3DL2 expression in the full biospecimen cohort (N=50). **E)** Representative flow cytometry histograms of KIR3DL2 staining including unstained, IgG2a κ isotype control, wild-type cells, KIR3DL2 knockout (KO) and KIR3DL2 knock-in in two CTCL cell lines with endogenous KIR3DL2 expression (HUT-78 and MAC-2A) and in one line lentivirally transduced with human KIR3DL2 (HH). nTPM, normalized transcripts per million; MFI, mean fluorescence intensity; D, donor; N, naïve (CD45RA+CD62L+); EM, early memory (CD45RA-CD62L-); CM, central memory (CD45RA-CD62L+); TEMRA, terminally differentiated effector memory (CD45RA+CD62L-). HSTL, hepatosplenic TCL; ENKTL, extranodal NK/TCL; PTCL, NOS, peripheral TCL, not otherwise specified; BIA-ALCL, breast implant–associated anaplastic large cell lymphoma; AITL, angioimmunoblastic TCL; pcγδTCL, primary cutaneous γδ TCL; nγδTCL, nodal γδ TCL; MEITL, monomorphic epitheliotropic intestinal TCL; T-LGL, T-cell large granular lymphocytic leukemia; ATLL, adult T-cell leukemia/lymphoma; MF-LCT, mycosis fungoides with large cell transformation; pcALCL, primary cutaneous anaplastic large cell lymphoma; T-PLL, T-cell prolymphocytic leukemia; MF, mycosis fungoides; nTFHL, nodal T-follicular helper cell lymphoma; ALCL, ALK+, anaplastic large cell lymphoma, ALK-positive; ALCL, ALK−, anaplastic large cell lymphoma, ALK-negative; EATL, enteropathy-associated TCL.

### Our proprietary anti-KIR3DL2 mAb is specific to KIR3DL2

An in-house mAb targeting KIR3DL2 was generated by hybridoma technology (Suppl. Methods). Hybridoma fusions were screened using flow cytometry on mouse fibroblast L cells transduced with human KIR3DL2 as well as TCL cell lines. This screening identified clone K36, which specifically bound to human KIR3DL2-transduced L cells and HUT-78 but not to parental L cells (**Suppl. Fig. 2A** and data not shown). Clone K36 selectively bound KIR3DL2-expressing HEK293T cells, with no cross reactivity to the other 14 KIR family members (**Suppl. Fig 2B**).

### Lentiviral delivery of 3DL2CAR constructs to primary T cells results in high and sustained transgene expression

We generated four clone K36-derived scFv-based 3DL2CAR constructs encoding different H/TM/CO and CD3ζ signaling domains. All encoded cdtBCMA, which served as a transduction marker and/or safety switch (**Fig 2A**). Lentiviral delivery of these constructs to activated primary T cells resulted in high and stable transduction efficiency as assessed by flow cytometry at day 3 and 7 after transduction (**Fig. 2B** and data not shown).

**Figure 2.**
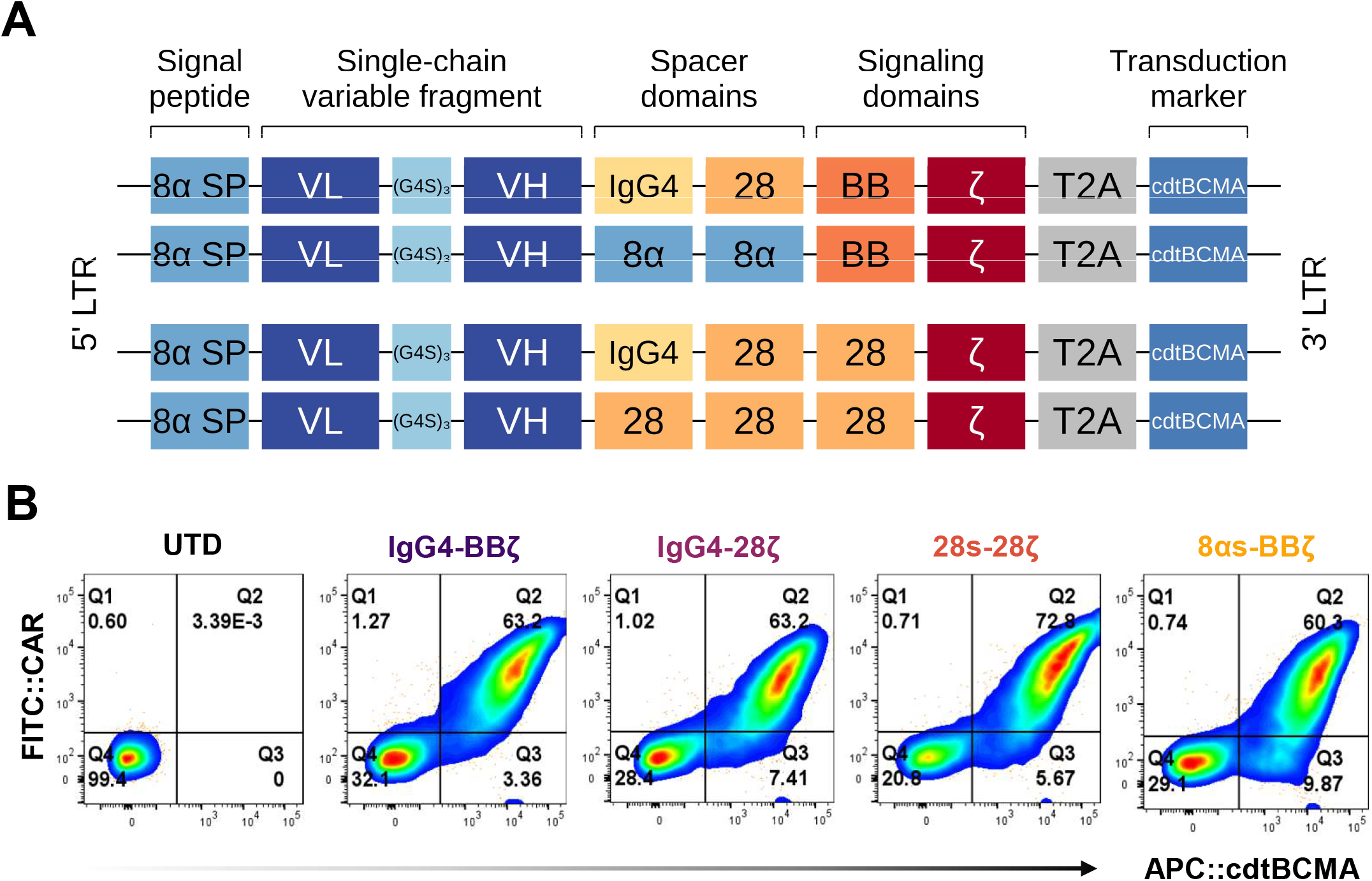
Lentiviral delivery of 3DL2CAR constructs to primary T cells results in high and sustained transgene expression. **A)** Schematic of 3DL2CAR constructs. Clone K36 VL and VH were joined by a (G₄S)₃ linker fused to IgG4+CH3, CD8α, CD28 hinge; CD28 or CD8α transmembrane and 4-1BB or CD28 costimulatory domains. All constructs encoded CD3ζ signaling domain and the transduction marker/safety switch cdtBCMA (a signaling-incompetent truncated form BCMA, lacking the intracellular domain but with two copies of the extracellular domain). **B)** 3DL2CAR constructs were lentivirally transduced into activated primary T cells. Transduction efficiency was estimated by flow cytometry on day 3 and day 7 post-transduction. Pseudocolor plots represent the binding of the anti-KIR3DL2 scFv to recombinant human KIR3DL2 protein with His tag as detected by anti-His antibody (ab) staining. cdtBCMA expression was detected by anti-BCMA ab staining. LTR, long terminal repeat; VL, variable light chain; VH, variable heavy chain; 28, CD28; 8α, CD8α; BB, 4-1BB; s, spacer.

### 3DL2CART cells mediate antigen-specific cytotoxicity, signaling, proliferation and cytokine release *in vitro*

We sought to assess the functional activity of 3DL2CART cells against non-target and target TCL cell lines in co-culture experiments. 3DL2CARs did not show cytotoxic activity against KIR3DL2 knockout (KO) HUT-78 or HH (KIR3DL2 negative), but exhibited high cytotoxic activity against HUT-78, HH^KIR3DL^^2^ and MAC-2A cell lines at effector to target (E:T) ratios as low as 1:10 and following repeated tumor challenges (**Fig. 3A** and **Suppl. Fig 3A**). Jurkat-Lucia™ NFAT were transduced with different 3DL2CARs and exhibited low tonic signaling and specific activation in response to HUT-78 and HH^KIR3DL^^2^, but not against HH (**Fig 3B** and Suppl. Methods). Moreover, 3DL2CART cells showed increased proliferation compared with donor-matched activated UTD T cells upon co-culture with KIR3DL2-expressing cells and to a lesser extent with antigen-negative isogenic clones. (**Fig. 3C** and **Suppl. Fig. 3B**). 3DL2CAR T-cells also exhibited potent cytokine production compared with control T cells when co-cultured with HUT-78 target cell line but not with KIR3DL2-negative HH (**Fig. 3D** and **Suppl. Fig 3C**).

**Figure 3.**
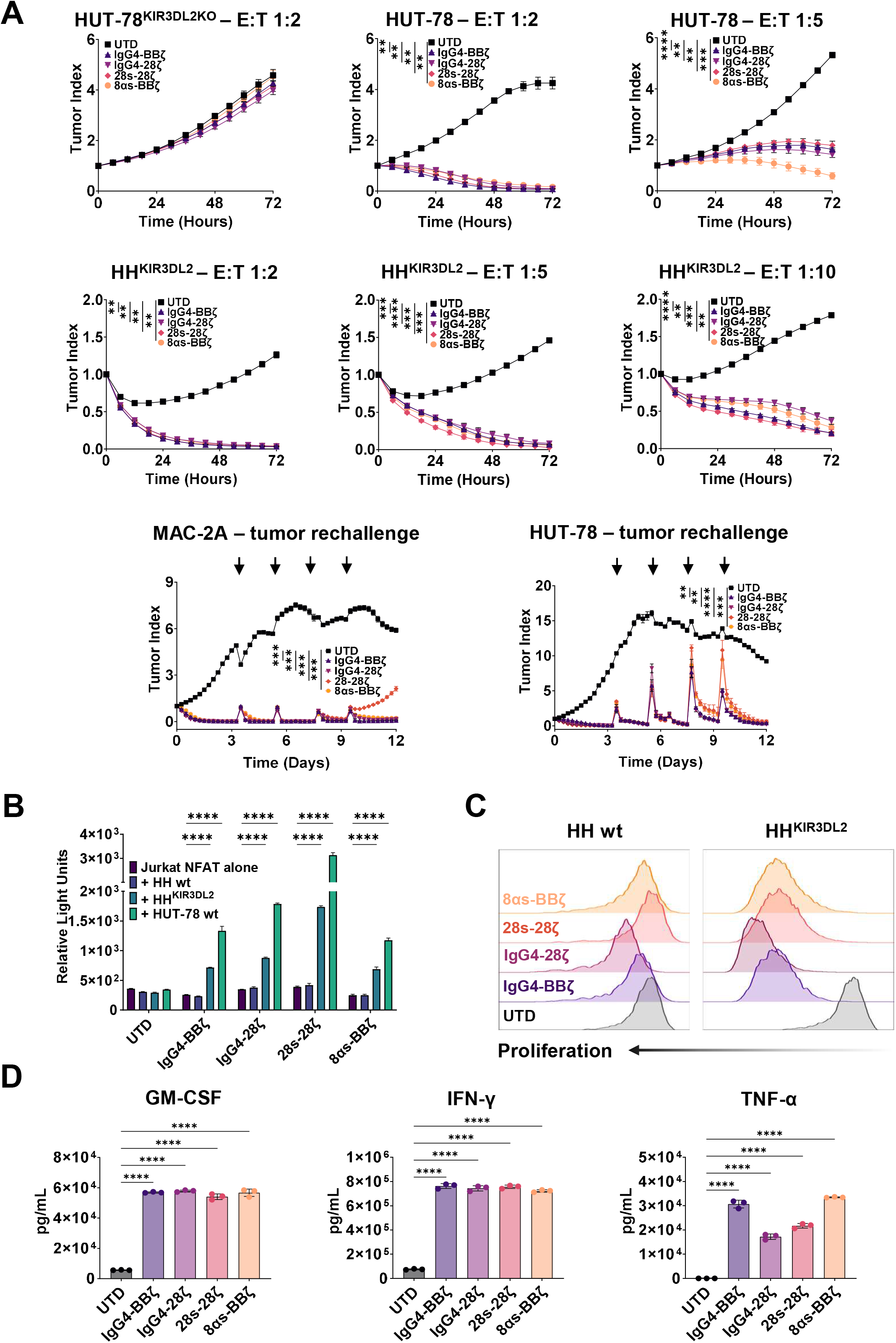
3DL2CAR T cells mediate antigen-specific cytotoxicity, signaling, proliferation and cytokine release *in vitro*. **A)** Isogenic KIR3DL2-knockout (KO) HUT-78, wild type HUT-78, KIR3DL2-expressing HH and wild type MAC-2A were lentivirally transduced to express eGFP and co-cultured in triplicate wells with or without expanded untransduced control T-cells or 3DL2CAR T-cells at the indicated effector to target ratios (E:T). Live tumor cell quantification was performed using Incucyte Live-Cell Imaging by monitoring eGFP^+^ cells. Mean ± SD of fold change in live tumor cells from baseline (Tumor Index) over time is shown for each condition. Vertical arrows denote re-challenge with fresh tumor cells. Statistical significance was determined by two-way ANOVA followed by Dunnett’s multiple comparisons test. Plots are representative of experiments performed with at least three independent donors. **B)** Jurkat-Lucia™ NFAT reporter cells were lentivirally transduced with 3DL2CAR constructs and co-cultured in triplicate wells with or without KIR3DL2^+^ or KIR3DL2^-^ TCL cell lines for 24 hours. Bar graphs represent mean ± SD of relative light units (RLU) measured in culture supernatants following incubation with QUANTI-Luc™. Statistical significance was determined by two-way ANOVA followed by Dunnett’s multiple comparisons test. **C)** 3DL2CAR or UTD T cells were labeled with CellTrace™ Far Red and co-cultured with antigen-negative or antigen-positive target cells for 96 hours. The proliferating fraction was estimated from the decrease in dye MFI on CD3⁺BCMA⁺ cells (CAR^+^ cells) or on CD3⁺ cells in the UTD group. Data are representative of three independent experiments with three different donors and three different target/non-target cell lines. **D)** 3DL2CAR and UTD T cells were co-cultured with HUT-78 cells at a 1:1 E:T ratio. Cytokine release was assessed in culture supernatants after 48 hours by multiplex assay (Meso Scale Discovery). Bar graphs show GM-CSF (left), IFN-γ (middle), and TNF-α (right), selected from a 10-cytokine panel. Statistical significance was determined by one-way ANOVA followed by Dunnett’s multiple comparisons test. Data are representative of two independent donors, assayed against both antigen-positive and antigen-negative cell lines. wt, wild type. Statistical significance was reported as: * for p < 0.05, ** for p < 0.01, *** for p < 0.001, and **** for p < 0.0001.

### 3DL2CART cells induce sustained tumor control and extend survival in two CTCL cell-line-derived xenograft models (CDX) and one HSTL PDX

We tested the *in vivo* anti-tumor activity of 3DL2CART cells in a CDX of HH^KIR3DL^^2^ (**Fig 4A**). All mice in all treatment groups exhibited rapid and complete tumor regression, except for one mouse in the 28s-28BBζ group that showed central nervous system involvement and was sacrificed due to progressive ataxia. Tumor burden was lower in all 3DL2CAR groups when compared with UTD-treated mice (**Fig. 4B**). Survival was also significantly increased in all 3DL2CAR-treated groups compared with UTD-treated group (**Fig. 4C**). Accordingly, there was a progressive weight increase in 3DL2CAR-treated mice (**Suppl. Fig. 4A**). Similarly, HUT-78-engrafted mice treated with IgG4-BBζ, IgG4-28ζ or 8αs-BBζ constructs demonstrated rapid tumor control, progressive weight gain, and prolonged survival (**Fig. 4D-F** and **Suppl. Fig. 4B**). Notably, constructs encoding 4-1BB costimulatory domain showed increased CAR T-cell expansion compared with IgG4-28ζ (**Fig. 4G**). We then tested IgG4-BBζ construct against CBTL-81777, a HSTL PDX model. CAR-treated mice showed increased survival and weight gain compared with control arms (**Fig. 5A, B and Suppl. Fig. 5A**). We delivered the same tumor and effector T cell dose using a different donor and sacrificed all mice 14 days after treatment administration. IgG4-BBζ treated-group exhibited robust expansion in peripheral blood and effectively controlled tumor growth in peripheral blood, spleen and liver (**Fig. 5C-E**).

**Figure 4.**
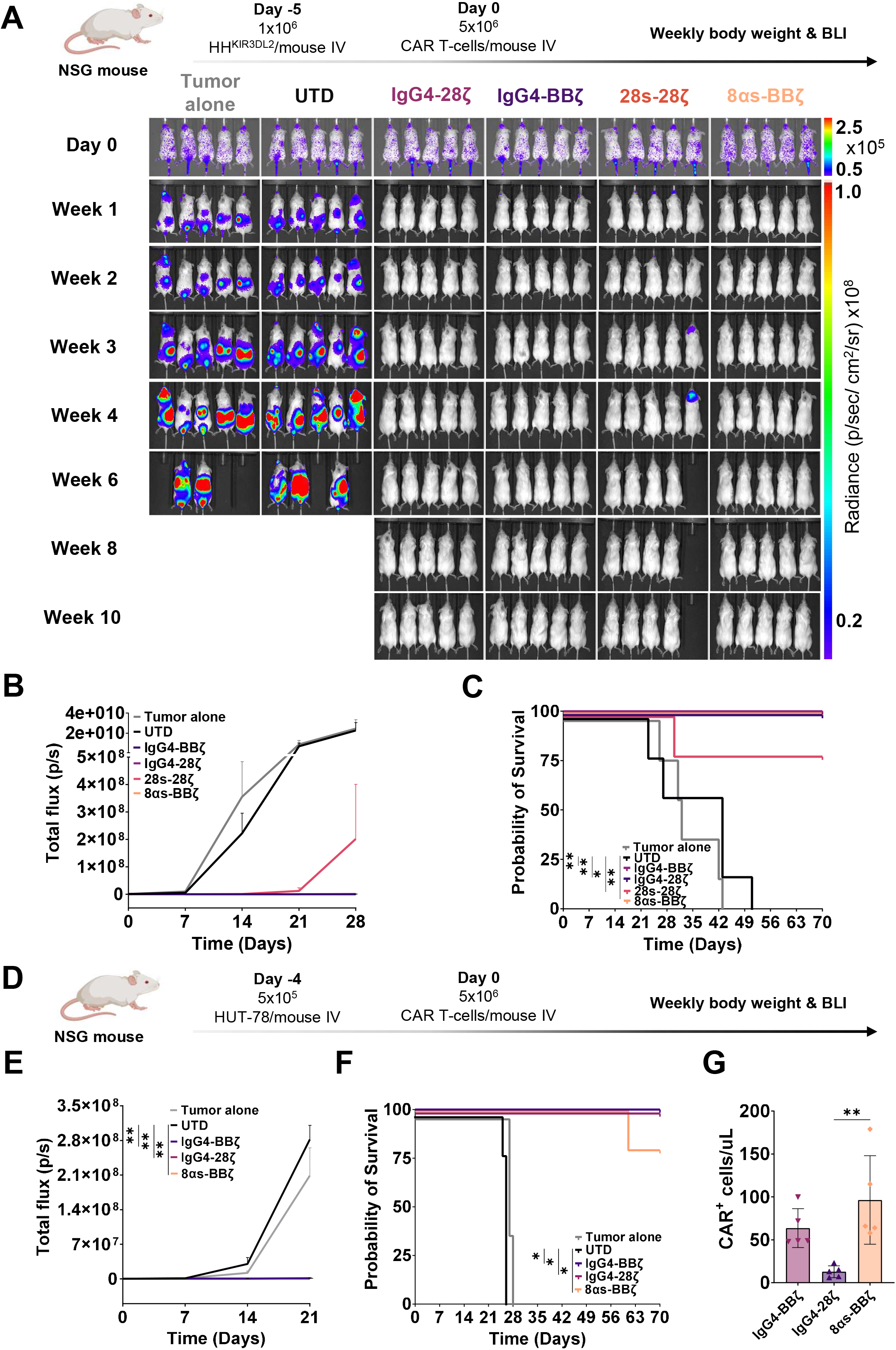
3DL2CAR T cells induce sustained tumor control and extend survival in two CTCL cell-line-derived xenograft models (CDX). **A**) Luciferase-labeled HH^KIR3DL^^2^ were injected via tail vein into NSG mice at doses of 1×10⁶ cells/mouse (day -5). On day 0, 5×10⁶ UTD or 3DL2CAR T cells were injected via tail vein. Tumor growth was monitored weekly by bioluminescence imaging (BLI). Created in BioRender. Aletti, F. M. (2026) https://BioRender.com/rkedlt1. **B)** HH^KIR3DL^^2^ tumor burden estimated by mean flux of photon emission for each group is shown at the indicated timepoints. Statistical significance was determined by two-way ANOVA followed by Dunnett’s multiple comparisons test. **C)** Kaplan–Meier survival curves of untreated, UTD or 3DL2CAR treated mice from the same experiment. Differences between each 3DL2CAR treatment group and the UTD control group were assessed using pairwise log-rank (Mantel–Cox) tests, with Holm–Bonferroni adjustment for four comparisons. Adjusted p values are shown. **D)** Luciferase-labeled HUT-78 were injected via tail vein into NSG mice at doses of 5×10^5^ cells/mouse (day -4). On day 5×10⁶ UTD or 3DL2CAR T cells were injected via tail vein. Tumor growth was monitored weekly by BLI. *Created in BioRender. Aletti, F. M. (2026)* https://BioRender.com/rkedlt1. **E)** HUT-78 tumor burden estimated by mean flux of photon emission for each group is shown at the indicated timepoints. Statistical significance was determined by two-way ANOVA followed by Dunnett’s multiple comparisons test. **F)** Kaplan–Meier survival curves of untreated, UTD or 3DL2CAR treated mice from the same experiment. Differences between each 3DL2CAR treatment group and the UTD control group were assessed using pairwise log-rank (Mantel–Cox) tests, with Holm–Bonferroni adjustment for three comparisons. Adjusted p values are shown. **G)** CAR^+^ cells (defined as live hCD45^+^HLA-A2^+^cdtBCMA^+^ cells) concentration in the peripheral blood on day 7 post treatment infusion. Statistical significance was determined by one-way ANOVA followed by Dunnett’s multiple comparisons test. Statistical significance was reported as: * for p < 0.05, ** for p < 0.01, *** for p < 0.001, and **** for p < 0.0001.

**Figure 5.**
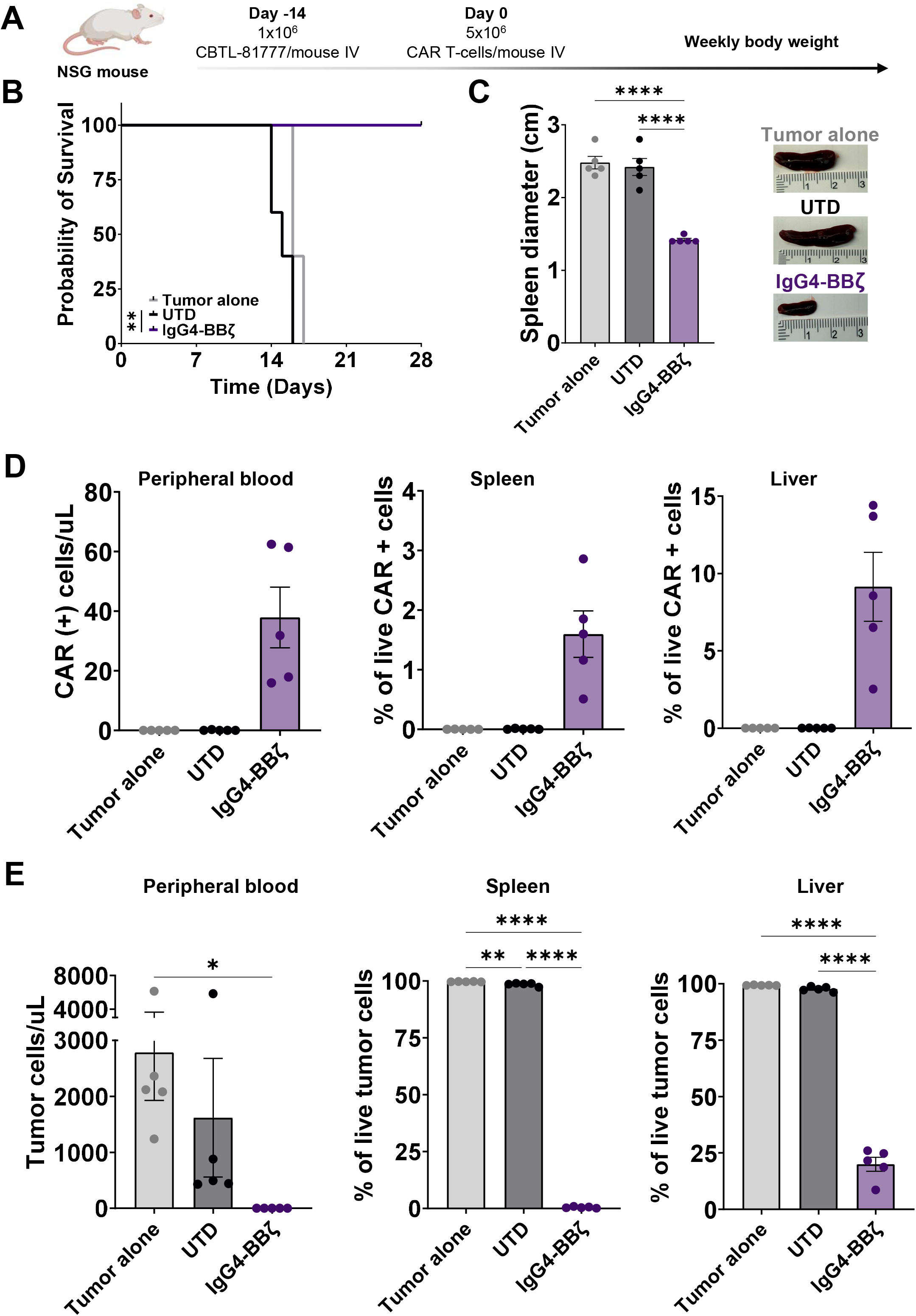
3DL2CAR T cells induce sustained tumor control and extend survival in a HSTL patient-derived xenograft models (PDX). **A)** CBTL-81777 cells were injected via tail vein into NSG mice at doses of 1×10⁶ cells/mouse (day -14). On day 0, 5×10⁶ UTD or IgG4.BBζ CAR T cells were injected via tail vein. *Created in BioRender. Aletti, F. M. (2026)* https://BioRender.com/rkedlt1. **B)** Kaplan–Meier survival curves of untreated, UTD or IgG4.BBζ treated mice from the same experiment. Differences between each IgG4.BBζ treatment group and the UTD were assessed using pairwise log-rank (Mantel–Cox) test. **C)** The same experimental setup was used with a different donor, and all mice were euthanized 2 weeks after treatment administration. The bar graph displays differences in the spleen interpolar diameter between groups. Statistical significance was determined by one-way ANOVA followed by Tukey’s multiple comparisons test. **D)** CAR^+^ cells (defined as live hCD45^+^HLA-A2^+^cdtBCMA^+^ cells) concentration in the peripheral blood (left) or CAR^+^ frequency among hCD45^+^ cells in spleen (middle) or liver (right) at necropsy. **E)** Tumor burden (defined as live hCD45^+^HLA-A2^-^) in peripheral blood (left), spleen (middle) or liver (right) in the same experiment. Statistical significance was determined by one-way ANOVA followed by Tukey’s multiple comparisons test. Statistical significance was reported as: * for p < 0.05, ** for p < 0.01, *** for p < 0.001, and **** for p < 0.0001.

### 3DL2CART cells show negligible off-target toxicity on healthy immune subsets and minimal fratricide

To screen for potential off-target activity of 3DL2CART cells, we co-cultured freshly thawed PBMCs with donor-matched activated UTD T cells, non-signaling 3DL2CART cells (3DL2CAR.Δ) and 3DL2CAR.BBζ T cells (**Fig. 6A**). No or minimal differences in NK and B cell counts were detected across all groups, whereas the total CAR-negative T cell fraction was increased in 3DL2CAR.BBζ, likely because of bystander proliferation related to CAR T-cell activation (**Fig. 6B** and **Suppl. Fig. 6A**). Indeed, when compared with 3DL2CAR.Δ, 3DL2CAR.BBζ T cells showed significantly increased proliferation suggesting antigen-dependent signaling (**Fig. 6C** and **Suppl. Fig. 6B**). We also did not detect KIR3DL2 upregulation in 19CAR and 3DL2CAR T cells upon repeated stimulation with respective target and non-target cells. (**Fig. 6D and Suppl. Fig. 6C**). We then tested our product for potential T-cell depletion and/or fratricide during manufacturing. First, we transduced donor-matched T cells with either eGFP or 3DL2CAR.Δ/3DL2CAR.BBζ and cocultured eGFP^+^ and CAR^+^ cells. Expansion rate of bystander eGFP^+^ cells was similar across all groups with no evidence of T-cell depletion in the signaling-competent CAR group when compared with control groups (**Fig. 6E**). There was also no significant difference in proliferation between signaling-competent and signaling-deficient CAR^+^ cells across three different donors, suggesting absence of significant fratricide (**Fig. 6F**). This was also substantiated by no difference in viable cells as assessed by Annexin V/Propidium iodide staining of co-cultures of UTD, 3DL2CARΔ or 3DL2CAR with donor-matched eGFP-labelled T cells (**Suppl. Fig. 6D** and Suppl. Methods).

**Figure 6.**
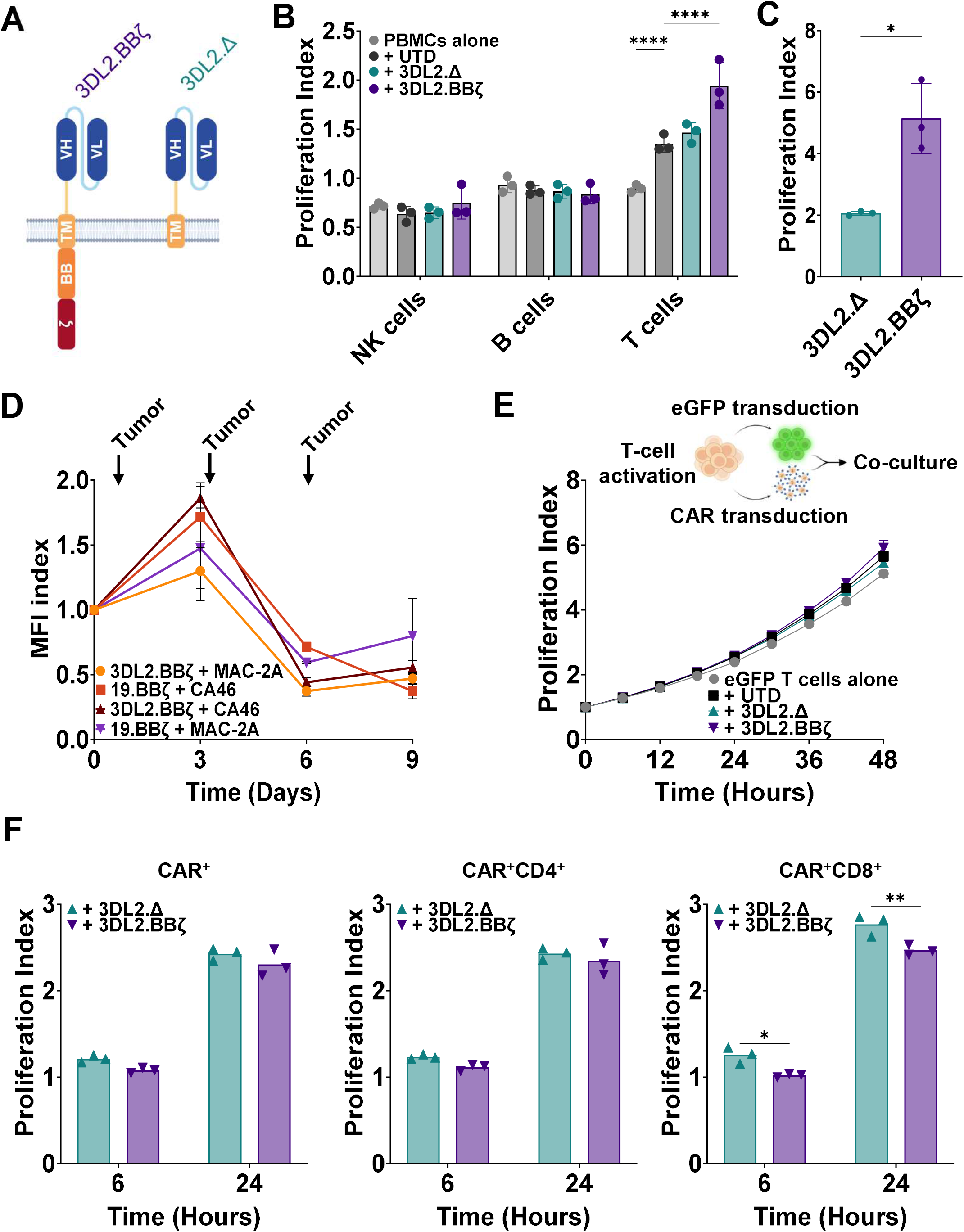
3DL2CAR T cells show negligible off-target toxicity on healthy immune subsets and minimal fratricide. **A)** Schematic of signaling-competent (3DL2.BBζ, left) and signaling-incompetent (3DL2.Δ, right) CAR constructs. Created in BioRender. Aletti, F.M. (2026) https://BioRender.com/qy9sj7b. **B)** Healthy donor PBMCs were co-cultured with or without UTD, 3DL2.Δ or 3DL2.BBζ for 48 hours at 1:5 E:T in triplicate wells. Bar graphs represent the mean ± SD of fold change in live cells from each subset normalized to baseline (proliferation index). Statistical significance was determined by two-way ANOVA followed by Dunnett’s multiple comparisons test. Data are representative of results obtained with three different donors. **C)** Proliferation index for 3DL2.Δ or 3DL2.BBζ CAR^+^ cells from the same experiment. Statistical significance was determined by an unpaired t test with Welch’s correction. **D)** 3DL2.BBζ or anti-CD19 (19.BBζ) CAR T cells were co-cultured in triplicate wells with CA46 (KIR3DL2^-^CD19^+^ Burkitt lymphoma cell line) or MAC-2A (KIR3DL2^+^CD19^-^) at a E:T of 1:1. A flat dose of 5×10^4^ tumor cells was added to each well on day 3, 6 and 9. The graph displays the fold MFI change of KIR3DL2 normalized to baseline MFI index) for each group at the indicated timepoints. Data are representative of results obtained with three different donors. Created in BioRender. Aletti, F.M. (2026) https://BioRender.com/xbmoqkp. **E)** eGFP-transduced primary T cells were incubated with or without donor-matched UTD, 3DL2.Δ or 3DL2.BBζ T cells for 48 hours at E:T of 1:1 in triplicate wells. Proliferation index was estimated using Incucyte Live-Cell Imaging by monitoring eGFP^+^ cells. Data are representative of results obtained with three different donors. **F)** Proliferation index values at 6 and 24 hours, assessed by flow cytometry on total CAR⁺, CD4^+^ CAR⁺ and CD8^+^CAR^+^ cells within the same experiment. Statistical significance was determined by two-way ANOVA followed by Šídák’s multiple comparisons test. Statistical significance was reported as: * for p < 0.05, ** for p < 0.01, *** for p < 0.001, and **** for p < 0.0001.

### Immunomagnetic depletion and safety-switch activation efficiently eliminate malignant cells spiked into healthy PBMCs

To mitigate the risk of malignant cell carryover and subsequent inadvertent transduction during autologous CAR T-cell manufacturing, we reproduced graded contamination of the apheresis product by spiking mCherry-tagged HUT-78 cells into healthy donor-derived PBMCs at low, intermediate and high ratios. Cell cultures were labeled with the anti-KIR3DL2 primary antibody followed by anti-fluorochrome magnetic microbeads and depleted on LD columns (**Fig. 7A** and Suppl. methods). At all three contamination levels, immunomagnetic depletion markedly reduced the mCherry⁺ fraction, with residual events confined to a KIR3DL2^dim/lo^ subset (**Fig. 7B**). As a second, independent layer of safety, we investigated whether malignant cells that escaped depletion and were inadvertently transduced could still be eliminated. HUT-78 cells were deliberately transduced with the 3DL2.BBζ construct, in which a ribosomal skipping element drives co-expression of cdtBCMA safety switch, and were co-cultured with healthy resting T cells in the presence of teclistamab, an anti-BCMA T-cell engager, or an isotype control (Suppl. Methods). As anticipated, surface KIR3DL2 MFI was modestly lower on CAR-transduced cells than on non-transduced controls, consistent with partial *cis*-masking of the target epitope by the CAR (**Fig. 7C**). Most importantly, exposure to subtherapeutic concentrations of teclistamab produced a significant reduction in HUT-78 BBζ cells relative to controls (**Fig. 7D**).

**Figure 7.**
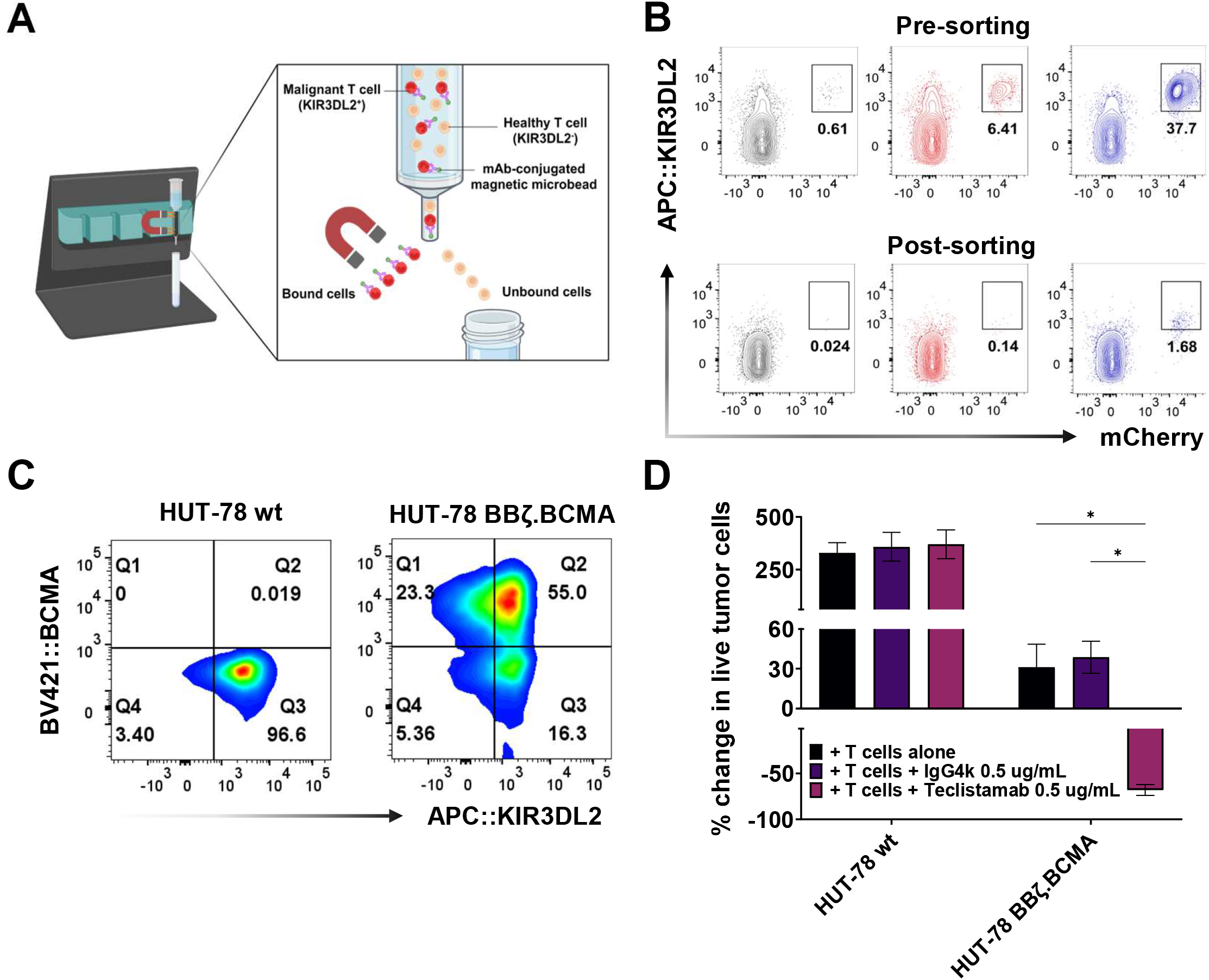
Immunomagnetic depletion and safety-switch activation efficiently eliminate malignant cells spiked into healthy PBMCs. **A)** mCherry-labeled HUT-78 cells were spiked at increasing concentrations into healthy donor PBMCs and incubated with an AF647-conjugated anti-KIR3DL2 antibody for 30 minutes. After washing, cells were labeled with anti-fluorophore magnetic microbeads and the suspension was passed through an LD column. Created in BioRender. Aletti, F. M. (2026) https://BioRender.com/ftimapd **B)** Flow cytometry contour plots of mCherry and KIR3DL2 expression in the input suspension (top) and in the flow-through fraction after LD column depletion (bottom). **C)** Flow cytometry plots of KIR3DL2 and BCMA expression in wild-type HUT-78 (left) and in HUT-78 transduced with the 3DL2CAR construct encoding the BCMA safety switch (HUT-78 BBζ.BCMA, right). **D)** HUT-78 WT and HUT-78 BBζ.BCMA cells were co-cultured with resting healthy donor T cells at an E:T ratio of 1:1, with or without IgG4κ isotype control or teclistamab at 0.5 µg/mL, in triplicate wells. Bar graphs show the percent change in the absolute count of live mCherry⁺ cells relative to baseline. Statistical significance was determined by one-way ANOVA followed by Dunnett’s multiple comparisons test. Statistical significance was reported as: * for p < 0.05, ** for p < 0.01, *** for p < 0.001, and **** for p < 0.0001.

## DISCUSSION

Here, we report KIR3DL2 as a promising target for CAR T-cell therapy of KIR3DL2-expressing subtypes of cutaneous and systemic TCLs. The restricted expression of KIR3DL2 in normal tissues and immune cells, coupled with its high expression across several TCL subtypes, including SS, transformed MF, ENKTL, HSTL, pc and nγδTCL, MEITL and others suggests that KIR3DL2 can be targeted safely with CAR T-cell therapy while minimizing on-target, off-tumor toxicity. Importantly, KIR3DL2 expression in defined NK and CD8^+^ TEMRA cells rather than the broader lymphoid compartment confers several potential advantages. First, any off-tumor activity would be limited to a subset of T and NK cells rather than the entire T-or NK-cell pool, substantially reducing the risk of profound immunosuppression and infectious complications. Second, fratricide during CAR T-cell manufacturing is expected to be minimal, eliminating the need for gene editing or other strategies to knockout or knockdown KIR3DL2 expression in the engineered T cells. Finally, in the autologous setting, KIR3DL2-expressing malignant cells can be selectively depleted from apheresis products prior to T-cell transduction, reducing the risk of inadvertent CAR transgene incorporation into tumor cells and the subsequent emergence of immune escape^30^.

In efficacy studies, we demonstrated that KIR3DL2-directed CAR T-cells are highly efficient at eliminating KIR3DL2^+^ tumor cells in a CAR-dependent and antigen-dependent manner. Importantly, they induced rapid tumor regression and significant prolongation of survival *in vivo* across antigen high and antigen low tumor xenografts including a PDX of HSTL. Together with the expected safety profile, these results support evaluation of this CAR T-cell product in patients with various KIR3DL2-expressing subtypes of cutaneous and systemic TCLs, a group with poor clinical outcomes. Current therapies including standard chemotherapy, targeted drugs, and stem cell transplantation often yield only transient responses and rarely achieve sustained remission^44^. Patients with SS, pcγδTCL and transformed MF represent the aggressive end of the CTCL spectrum and carry poor prognoses, with reported 5-year disease-specific survival of 36% and 11% for SS and pcγδTCL respectively^45^, and median overall survival <20 months after large-cell transformation of MF^46^. MEITL is an aggressive subtype of intestinal TCL with 1-year survival rates of 30% to 40%^42^; HSTL also carries a similarly poor prognosis, with long-term remission essentially confined to patients consolidated with allogeneic transplant^42^. Outcomes for localized ENKTL have improved substantially with radiotherapy and non-anthracycline-based regimens; however, advanced-stage disease retains a poor prognosis, with 5-year overall survival below 40%^47^. More generally, with the exception of ALCL which carries a more favorable prognosis^48,49^, and large granular lymphocytic leukemia which follows a generally indolent course^50^, TCL subtypes herein described typically show poor or short-lived responses to first-line therapy and available salvage options rarely produce durable remissions^51^. Thus, a KIR3DL2-directed CAR T-cell product would offer a novel therapeutic strategy for these patients with major unmet clinical need.

To date, most CAR T-cell approaches for T-cell malignancies have focused on targeting pan-T-cell antigens^17,19,52,53^, with CD5- and CD7-directed products representing the most clinically advanced strategies^18,22,24,54^. Although these therapies have demonstrated high response rates, they are frequently associated with prolonged T-cell aplasia, infectious complications^20,21,54,55^, and substantial non-relapse mortality, particularly among patients who do not proceed to consolidative allogeneic hematopoietic stem cell transplantation^24^. In addition, overcoming fratricide during manufacturing has required complex engineering approaches, including gene-editing and antigen-silencing strategies for both CD5- and CD7-targeted CAR T-cell products^20,22,54^. By contrast, our KIR3DL2-directed CAR T-cell platform largely avoids these challenges by targeting an antigen with highly restricted expression in normal lymphocyte subsets, thereby minimizing both fratricide and broad immune depletion while simplifying manufacturing. Indeed, we did not detect any significant NK, T, and B-cell depletion by 3DL2CART cells in co-culture studies with donor-matched PBMCs. We also did not observe evidence of fratricide nor antigen upregulation on CAR T cells during manufacturing or after serial stimulation with tumor cells.

The safety of KIR3DL2 targeting is further supported by clinical experience with the KIR3DL2-directed monoclonal antibody lacutamab. Across multiple clinical studies, lacutamab demonstrated a favorable safety profile together with promising antitumor activity in patients with r/r CTCL including SS^56–59^. These findings have established KIR3DL2 as a clinically tractable target in CTCL and potentially systemic TCLs. The key remaining question is therefore not whether KIR3DL2 can be effectively targeted, but rather which therapeutic modality can most effectively exploit this target. Drawing on the experience of CAR T-cell therapy in B-cell malignancies^60,61^, and supported by mechanistic considerations, we reasoned that KIR3DL2-directed CAR T cells may offer important advantages over unconjugated antibodies targeting the same antigen. First, CAR T cells mediate direct cytotoxicity independent of endogenous effector-cell function, whereas monoclonal antibodies rely primarily on antibody-dependent cellular cytotoxicity and antibody-dependent cellular phagocytosis. Second, CAR T cells can be engineered to recognize and eliminate tumor cells expressing low levels of target antigen^62^, potentially maintaining activity in settings where antibody-mediated effector mechanisms are suboptimal^63,64^. Finally, as living drugs capable of *in vivo* expansion and long-term persistence, CAR T cells are not constrained by the pharmacokinetic limitations of repeated antibody administration, potentially enabling more durable antitumor responses.

We recognize that the development of an autologous CAR T-cell product for TCLs that may manifest with a leukemic phase raises important safety considerations, particularly the risk of inadvertent transduction of malignant cells during manufacturing. While an allogeneic CAR T-cell approach may eliminate this risk completely^65,66^, we demonstrate that this could be overcome in the autologous setting by incorporating two complementary and independent safeguards. First, we showed that KIR3DL2^+^ cells can be depleted prior to T-cell transduction, thereby reducing the malignant cell burden entering the manufacturing process. Second, we incorporated a truncated BCMA safety switch within the same expression cassette of our CAR construct, based on an approach previously developed by our group^43^. This built-in strategy enables selective targeting and elimination of any transduced malignant clone that might emerge by using an approved BCMA-directed therapeutic agent, such as teclistamab. Together, these layered safeguards are designed to minimize the risk of malignant cell transduction while providing a clinically actionable mechanism for intervention should such an event occur.

In conclusion, our study establishes that KIR3DL2-directed CAR T-cell therapy is feasible and offers several potential safety advantages over strategies targeting pan-T-cell antigens. These findings provide the rationale for first-in-human evaluation of this approach, which would offer a transforming therapeutic alternative for patients with KIR3DL2-expressing cutaneous and systemic TCLs.

## Acknowledgements

This study was supported by the University of Texas MD Anderson Cancer Center T/NK-cell Malignancies Moonshot (SSN and FV), Lymphoma Research Foundation Postdoctoral Fellowship Grant (TRP 6591-20, FMA), Binaytara (Shaji Kumar Hematology Research Award, FMA) and National Institutes of Health NCI Cancer Center Support Grant to the University of Texas MD Anderson Cancer Center (P30 CA016672) (Flow Cytometry and Cellular Imaging Facility). The MD Anderson Lymphoma Tissue Bank is supported by KW Cares Foundation.

## Authorship Contributions

FMA generated and screened the hybridomas, performed the antibody and CAR subcloning, designed and performed the *in vitro* and *in vivo* experiments, analyzed the data, prepared the figures, and wrote the manuscript; JW generated and screened hybridomas, helped with antibody and CAR subcloning, designed the *in vitro* and *in vivo* experiments; CCL designed the *in vitro* and *in vivo* experiments, contributed to the development of BCMA safety switch, analyzed the data, optimized CBTL-81777 PDX dosing and functional assays protocols; JL generated the lentiviral plasmid pLVMG, developed the BCMA safety switch and designed the *in vitro* and *in vivo* experiments; FD supervised and performed data analysis; AR designed and supported *in vitro* and *in vivo* experiments and optimized HH CDX dosing; MDR designed and performed PBMCs phenotyping by flow cytometry, analyzed data and designed the *in vitro* and *in vivo* experiments; SK performed *in vitro* and *in vivo* experiments; XC helped sourcing all the reagents and supported experiments; SP, YT, C-YL contributed to the development and optimization of lentiviral vector and BCMA safety switch and supported the experiments; J.C. and F.C., designed and supported *in vitro* and *in vivo* experiments; KS, SDR, SGR, SM, JW supported *in vitro* and *in vivo* experiments; KC and NS provided guidance on *in vitro* and *in vivo* experiments; CK performed IHC staining of the patient biospecimens; MRG and LTB generated a MF-LCT PDX included in the IHC staining; SPI contributed to the study rationale; FV and NJD analyzed IHC data; SSN conceived the study, designed the *in vitro* and *in vivo* experiments, analyzed data, obtained funding and wrote the manuscript. All authors reviewed, provided critical feedback, and approved the final manuscript.

