## Supplementary figure legends & methods for "A novel KIR3DL2-directed CAR T-cell therapy for cutaneous and systemic T-cell lymphomas"

**SUPPLEMENTARY MATERIAL**

**SUPPLEMENTARY FIGURES**

**Supplementary figure 1. KIR3DL2 is expressed across different TCL patient biospecimens and cell lines. A)** Additional IHC pictures for KIR3DL2 in patient biospecimens of ALCL, ALK+ (KIR3DL2$^{-}$) and ENKTL, MEITL (KIR3DL2-positive). **B)** KIR3DL2 antigen density of cell lines and HSTL PDX (CBTL-81777) tested in this study, expressed as antibody binding capacity (molecules/cell). ALCL, ALK+, anaplastic large cell lymphoma, ALK-positive; HSTL, hepatosplenic TCL; ENKTL, extranodal NK/TCL; MEITL, monomorphic epitheliotropic intestinal TCL.

**Supplementary figure 2. Our proprietary anti-KIR3DL2 mAb is specific to KIR3DL2. A)** Hybridoma supernatants were incubated with HUT-78 cells and stained with an AF647-conjugated goat anti-mouse IgG secondary antibody. Dot plots obtained by flow cytometry show HUT-78 cells stained with secondary antibody alone (left) or with clone K36 supernatant (right). **B)** HEK293T cells were transfected with each of the 15 human KIR family members, and binding of clone K36 was assessed by flow cytometry. Histograms show MFI values for each KIR tested. KIR, killer-cell immunoglobulin-like receptor.

**Supplementary figure 3. Reproducibility of *in vitro* functional assays across different donors. A)** HUT-78, KIR3DL2-expressing HH and MAC-2A were lentivirally transduced to express eGFP or mCherry and co-cultured in triplicate wells with or without expanded untransduced control T-cells or 3DL2CAR T-cells at the indicated effector to target ratios (E:T). Live tumor cell quantification was performed using Incucyte Live-Cell Imaging by monitoring eGFP/mCherry$^{+}$ cells. Mean ± SD of fold change in live tumor cells from baseline (Tumor Index) over time is shown for each condition. Statistical significance was determined by two-way ANOVA followed by Dunnett's multiple comparisons test. **B)** 3DL2CAR or UTD T cells were labeled with CellTrace™ Far Red and co-cultured with antigen-negative or antigen-positive target cells for 96 hours. Bar graphs represent the fold change in CellTrace™ Far Red MFI normalized to baseline (MFI index). Far Red MFI was estimated on CD3$^{+}$BCMA$^{+}$ cells (CAR$^{+}$ cells) or on CD3$^{+}$ cells in the UTD group. Statistical significance was determined by two-way ANOVA followed by Dunnett's multiple comparisons test. **C)** 3DL2CAR and UTD T cells were co-cultured with HH or HUT-78 cells at a 1:1 E:T ratio. Cytokine release was assessed in culture supernatants after 48 hours by multiplex assay (Meso Scale Discovery). Bar graphs show GM-CSF, IFN-γ, and TNF-α, IL-2 selected from a 10-cytokine panel. Statistical significance was determined by two-way ANOVA followed by Dunnett's multiple comparisons test. Statistical significance was reported as: * for $p < 0.05$, ** for $p < 0.01$, *** for $p < 0.001$, and **** for $p < 0.0001$.

**Supplementary figure 4. Bioluminescence imaging of tumor burden and body weight monitoring in KIR3DL2$^+$ CDXs. A)** Luciferase-labeled HH$^{KIR3DL2}$ were injected via tail vein into NSG mice at doses of $1\times10^6$ cells/mouse (day -5). On day 0, $5\times10^6$ UTD or 3DL2CAR T cells were injected via tail vein. The graph shows the mean ± SD of percent change in body weight for each group relative to baseline (day −5). *Created in BioRender. Aletti, F. M. (2026) https://BioRender.com/rkedlt1.* **B)** Luciferase-labeled HUT-78 were injected via tail vein into NSG mice at doses of $5\times10^5$ cells/mouse (day -4). On day $5\times10^6$ UTD or 3DL2CAR T cells were injected via tail vein. Tumor growth and body weight were monitored weekly. *Created in BioRender. Aletti, F. M. (2026) https://BioRender.com/rkedlt1.* **C)** Mean ± SD of the percent change in body weight relative to baseline (day −4) for each group over time is shown.

**Supplementary figure 5. Body weight monitoring in CBTL-81777 xenograft. A)** CBTL-81777 cells were injected via tail vein into NSG mice at doses of $1\times10^6$ cells/mouse (day -14). On day 0, $5\times10^6$ UTD or IgG4.BBζ CAR T cells were injected via tail vein. The graph shows the mean ± SD of percent change in body weight for each group relative to baseline (day −5).

**Supplementary figure 6. Complementary analyses of on-target toxicity and fratricide. A)** Healthy donor PBMCs from two additional donors were co-cultured with or without UTD, 3DL2.Δ or 3DL2.BBζ for 48 hours at 1:5 E:T in triplicate wells. Bar graphs represent the mean ± SD of fold change in live cells from each subset normalized to baseline (proliferation index). Statistical significance was determined by two-way ANOVA followed by Tukey's multiple comparisons test. Data are representative of results obtained with three different donors. **B)** Proliferation index for 3DL2.Δ or 3DL2.BBζ CAR$^+$ cells from the same experiment. Statistical significance was determined by an unpaired t test with

Welch's correction. **C)** 3DL2.BBζ or anti-CD19 (19.BBζ) CAR T cells from two additional donors were co-cultured in triplicate wells with CA46 (KIR3DL2$^{-}$CD19$^{+}$ Burkitt lymphoma cell line) or MAC-2A (KIR3DL2$^{+}$CD19$^{-}$) at a E:T of 1:1. A flat dose of $5x10^4$ tumor cells was added to each well on day 3, 6 and 9. The graph displays the fold MFI change of KIR3DL2 normalized to baseline (MFI index) for each group at the indicated timepoints. **E)** Apoptosis was assessed by Annexin V and propidium iodide (PI) staining. The fractions of viable (Annexin V$^{-}$PI$^{-}$), early apoptotic (Annexin V$^{+}$PI$^{-}$), late apoptotic (Annexin V$^{+}$PI$^{+}$), and necrotic (Annexin V$^{-}$PI$^{+}$) cells within the whole culture are shown. Bars represent mean ± SD of triplicate wells at baseline and 24 hours from one representative experiment. Statistical significance was reported as: * for $p < 0.05$, ** for $p < 0.01$, *** for $p < 0.001$, and **** for $p < 0.0001$. PI, propidium iodide.

### SUPPLEMENTARY METHODS

#### Immunohistochemistry studies

Immunohistochemical staining for KIR3DL2 was performed using a polyclonal anti-KIR3DL2 antibody (Thermo Fisher Scientific) at a dilution of 1:100. All tissue sections underwent heat-induced epitope retrieval in Tris-EDTA buffer (pH 9.0). Endogenous peroxidase activity was blocked with 3% hydrogen peroxide ($H_2O_2$) for 30 minutes. To minimize nonspecific antibody binding, a serum-free blocking solution (Dako) was applied for 30 minutes at room temperature. Tissue sections were then incubated with the primary antibody overnight at 4 °C. Immunoreactivity was visualized using 3,3′-diaminobenzidine (DAB) as the chromogen. HUT-78 cells (SS cell line) and cellular block from CBTL-81777, a HSTL PDX, were used as positive controls. KIR3DL2 expression was assessed according to the percentage of positive cells, recorded in 10% increments, and staining intensity, graded as negative (0), weak (1+), moderate (2+), or strong (3+). A final H-score (HS) was calculated as follows: HS = [(% of cells with 1+ staining) × 1] + [(% of cells with 2+ staining) × 2] + [(% of cells with 3+ staining) × 3], yielding a score ranging from 0 to 300.

#### Anti-KIR3DL2 mAb production

L cells (mouse fibroblast cell line) were lentivirally transduced with human KIR3DL2 and injected into the footpad of BALB/c mice twice weekly for a total of six weeks. Mice were euthanized and B cells were harvested from popliteal lymph nodes and fused with Sp2/0-Ag14 cells (murine myeloma cell line) using the ClonaCell™-HY Hybridoma Generation Kit (STEMCELL Technologies) according to the manufacturer's protocol. Hybridomas were cultured in a methylcellulose-based semi-solid selective medium for 14 days, then transferred to liquid medium and expanded for a further week. Clone supernatants were screened by flow cytometry against parental L cells, KIR3DL2-transduced L cells, HUT-78, and SU-DHL-1 (a KIR3DL2-negative TCL cell line), using AF647-conjugated goat anti-mouse IgG (Jackson ImmunoResearch) as secondary antibody, leading to the identification of clone K36. Variable region sequences were obtained by RT-PCR using SMARTer mouse BCR kit (Takara Bio) according to the manufacturer's instructions. The resulting amplicons were cloned into a plasmid by TOPO™-TA™ cloning (Invitrogen) and sequenced by Sanger sequencing. The VH and VL regions were synthesized as eBlocks (IDT) and cloned into a lentiviral vector.

**Antibody purification and conjugation**

Hybridoma culture supernatant was clarified by centrifugation, concentrated on Pierce Protein Concentrators PES, 30K MWCO (Thermo Fisher Scientific), and the IgG fraction was affinity-purified using the NAb Protein G Spin Kit (Thermo Fisher Scientific) according to the manufacturer's instructions. Eluted antibody was neutralized and dialyzed against amine-free PBS at 4°C overnight using Slide-A-Lyzer Dialysis Devices, 20K MWCO (Thermo Fisher Scientific). Antibody concentration was determined by absorbance at 280 nm. Purified K36 was then conjugated to Alexa Fluor 647 (Invitrogen) following the manufacturer's protocol, and unbound dye was removed with the spin columns supplied with the kit.

**Assessment of K36 binding specificity across the human KIR family**

Coding sequences for each human KIR protein were cloned into the pcDNA3.1 vector (Addgene) upstream of a 2A peptide and eGFP, used as a transfection marker. HEK293T cells were transfected with Lipofectamine 3000 and cultured for 48 hours. Cells were then harvested by trypsinization, resuspended, and stained with AF647-conjugated K36 antibody. Samples were acquired by flow cytometry, and binding specificity was assessed as MFI relative to the negative control within the eGFP$^+$ population.

**Jurkat NFAT reporter luciferase release assay**

Jurkat-Lucia™ NFAT reporter cells were lentivirally transduced with 3DL2CAR constructs and expanded for 5 days. To obtain a pure population, cdtBCMA$^+$ cells were sorted and expanded for a further 3 days. CAR$^+$ cells were then co-cultured with KIR3DL2$^+$ or KIR3DL2$^-$ cell lines at an E:T ratio of 1:1 for up to 48 hours. Twenty microliters of supernatant were transferred to an opaque black 96-well plate, and 50 µL of QUANTI-Luc™ 4 Lucia/Gaussia substrate (InvivoGen), prepared according to the manufacturer's instructions, were added to each well. Bioluminescence was measured on a VICTOR Nivo multimode plate reader (PerkinElmer).

**Off-target and fratricide studies**

Activated untransduced (UTD) T cells, non-signaling CAR T cells, and signaling-competent CAR T cells were co-cultured with donor-matched peripheral blood mononuclear cells (PBMCs) at an effector-to-target (E:T) ratio of 1:5 in 96-well plates for 48 hours. Following co-culture, cells were stained and analyzed by flow cytometry. The proliferation index was calculated as the fold change in absolute viable cell counts relative to baseline for each cell subset. Absolute cell counts were determined by adding 10 µL of CountBright™ Absolute Counting Beads (Thermo Fisher Scientific) to each sample and applying the following formula: absolute cell count = (number of cell events / number of bead events) × total number of beads added. Cell concentrations (cells/mL) were calculated by dividing the absolute cell count by the initial sample volume (mL), accounting for dilution factors when applicable. A minimum of 1,000 bead events was acquired per sample. To evaluate KIR3DL2 expression following repeated antigen stimulation, an in-house-generated anti-CD19 CAR construct containing an FMC63-derived single-chain variable fragment (scFv), a CD8α spacer, and a 4-1BB costimulatory domain was used as a control. Anti-KIR3DL2 and anti-CD19 CAR T cells were co-cultured with target or non-target cells at an E:T ratio of 1:1 for 9 days. Fresh tumor cells were added on days 3 and 6 at the same dose used at the start of the co-culture. KIR3DL2 expression was assessed by flow cytometry at baseline and on days 3, 6, and 9, and reported as mean fluorescence intensity (MFI) within the cdtBCMA-positive T-cell population. For bystander T-cell toxicity and fratricide assays, T cells were isolated and activated as previously described. T cells were transduced with either eGFP or signaling-competent or signaling-incompetent CAR constructs and expanded until day 3 post-activation. At this time point, $eGFP^+$ T cells were co-cultured with UTD or CAR T cells for 48 hours. The expansion of $eGFP^+$ T cells were monitored using the Incucyte Live-Cell Analysis System (Sartorius), with images acquired every 6 hours. Changes in $eGFP^+$ T-cell numbers were assessed by measuring the mean green area confluence over time and expressed as fold change relative to baseline. CAR T-cell expansion was assessed at 6 and 24 hours by flow cytometry using absolute cell counting.

For viability assays cells were harvested, washed once in PBS and once in 1X binding buffer (Invitrogen), and resuspended in 1× binding buffer at 1–5 × $10^6$ cells/mL. Annexin V conjugate (Invitrogen) was added, and samples were incubated for 10–15 minutes at room temperature, protected from light per manufacturer's protocol. Cells were washed in 1X binding buffer, resuspended in 200 µL, and 5 µL of propidium iodide (Invitrogen) staining solution were added immediately before acquisition. Samples were analyzed by flow cytometry.

**Malignant cell depletion and safety switch activation**

Ten million PBMCs from healthy donors were spiked with increasing numbers of mCherry-tagged HUT-78 cells and incubated with AF647-conjugated anti-KIR3DL2 (clone K36) for 30 min at room temperature. After washing, cells were labeled with anti-PE-Cy5/AF647 MicroBeads (Miltenyi Biotec) according to the manufacturer's instructions and depleted on LD columns (Miltenyi Biotec). The flow-through fraction was analyzed by flow cytometry for the presence of residual $mCherry^+$ cells. For safety switch experiments, $mCherry^+$ HUT-78 cells were lentivirally transduced with the anti-KIR3DL2 CAR. After 72 h, transduced cells were co-cultured with isolated T cells at an E:T ratio of 1:1, with or without the addition of IgG4κ isotype control or teclistamab at the indicated concentrations. Percentage change in live cells normalized to baseline was determined by bead-based absolute cell counting through flow cytometry analysis.
