## Supplementary figures for "A novel KIR3DL2-directed CAR T-cell therapy for cutaneous and systemic T-cell lymphomas"

A

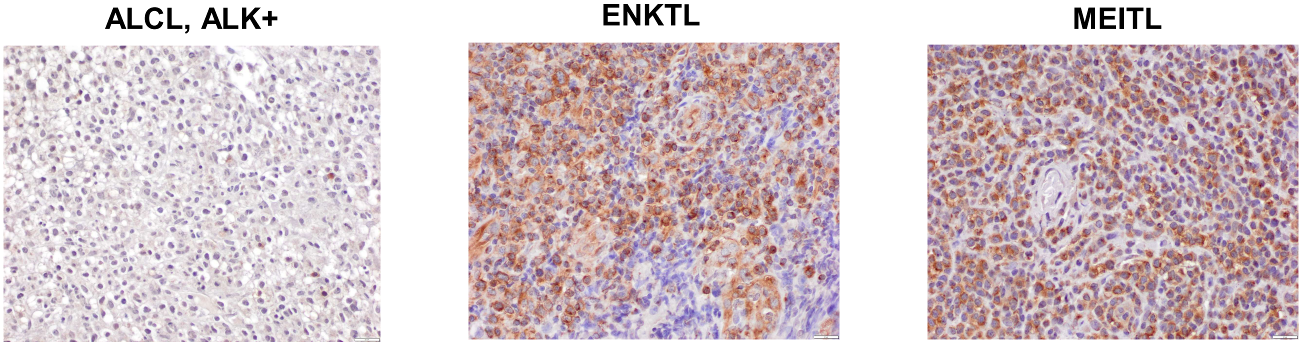


B

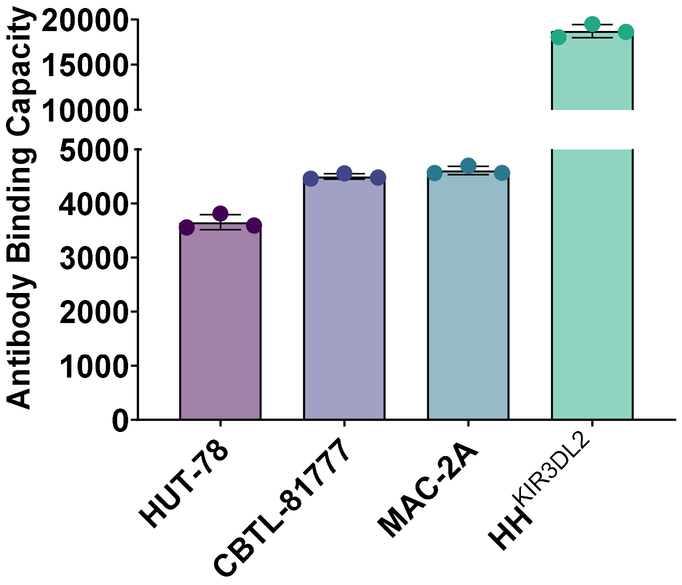


Suppl. Fig. 1

A

B

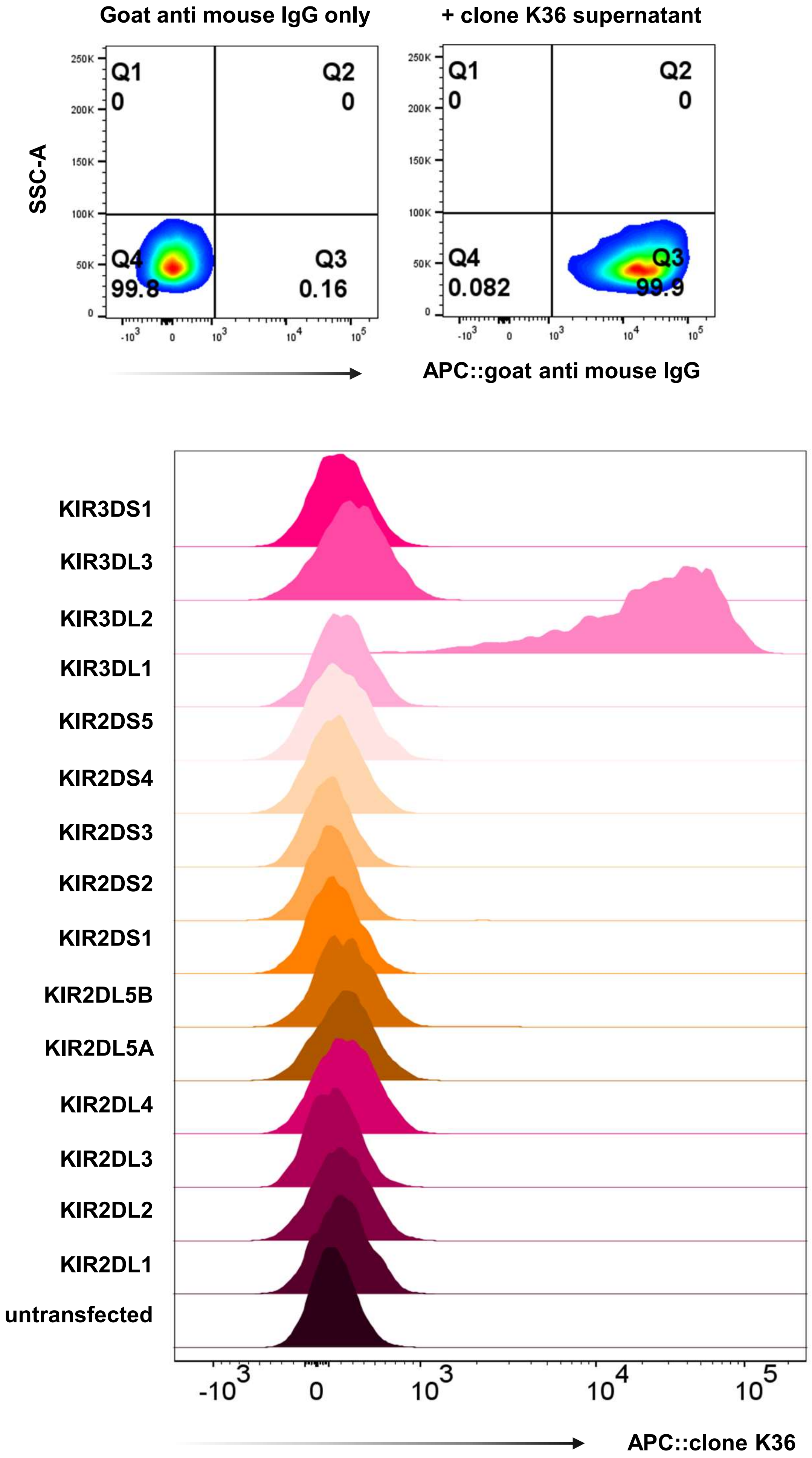


Suppl. Fig. 2

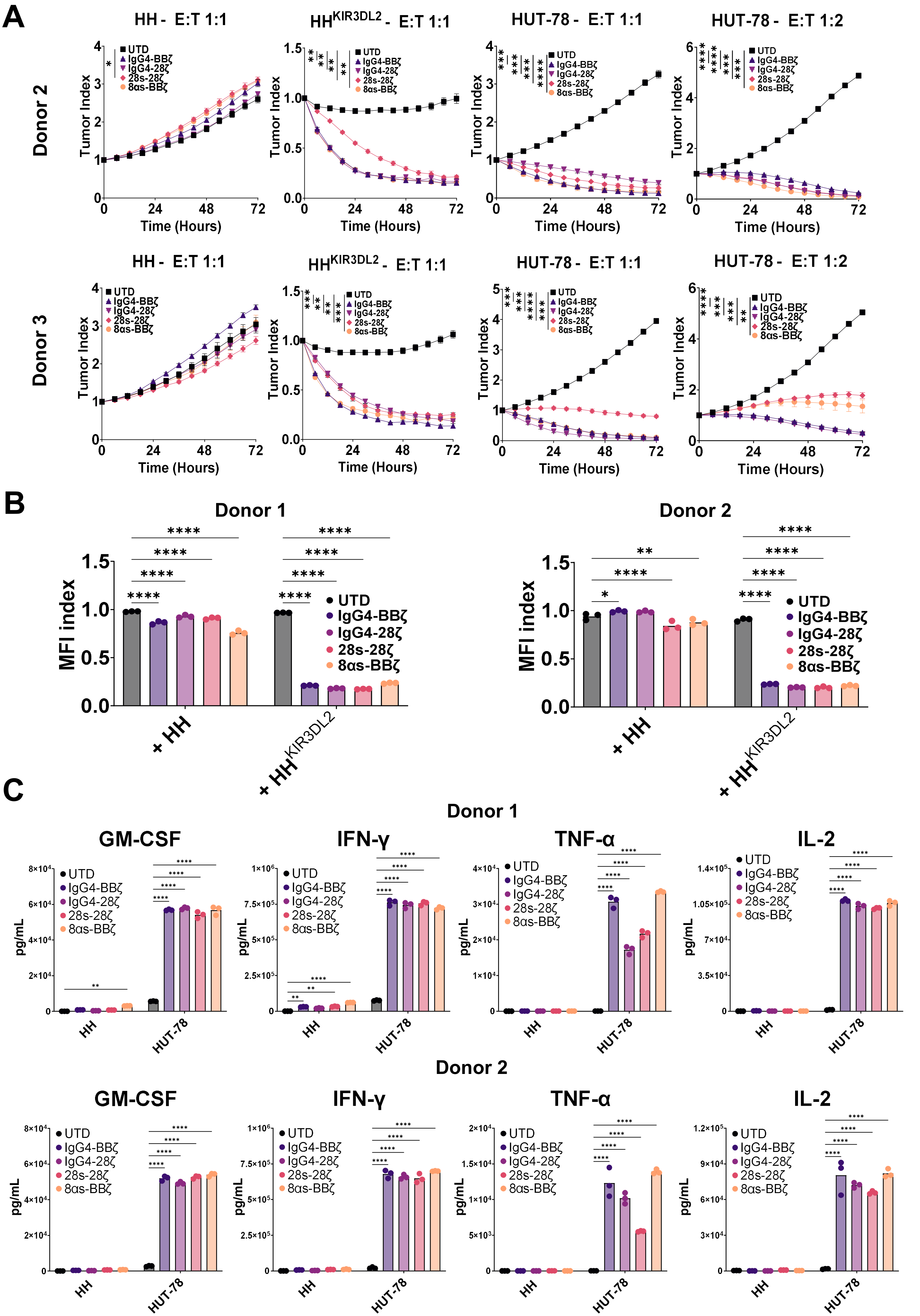
A
HH - E:T 1:1
HHKIR3DL2 - E:T 1:1
HUT-78 - E:T 1:1
HUT-78 - E:T 1:2
Donor 2
Donor 3
Tumor Index
Time (Hours)
UTD
IgG4-BBζ
IgG4-28ζ
28s-28ζ
8αs-BBζ
B
Donor 1
Donor 2
MFI index
+ HH
+ HHKIR3DL2
C
Donor 1
Donor 2
GM-CSF
IFN-γ
TNF-α
IL-2
pg/mL
HH
HUT-78

Suppl. Fig. 3

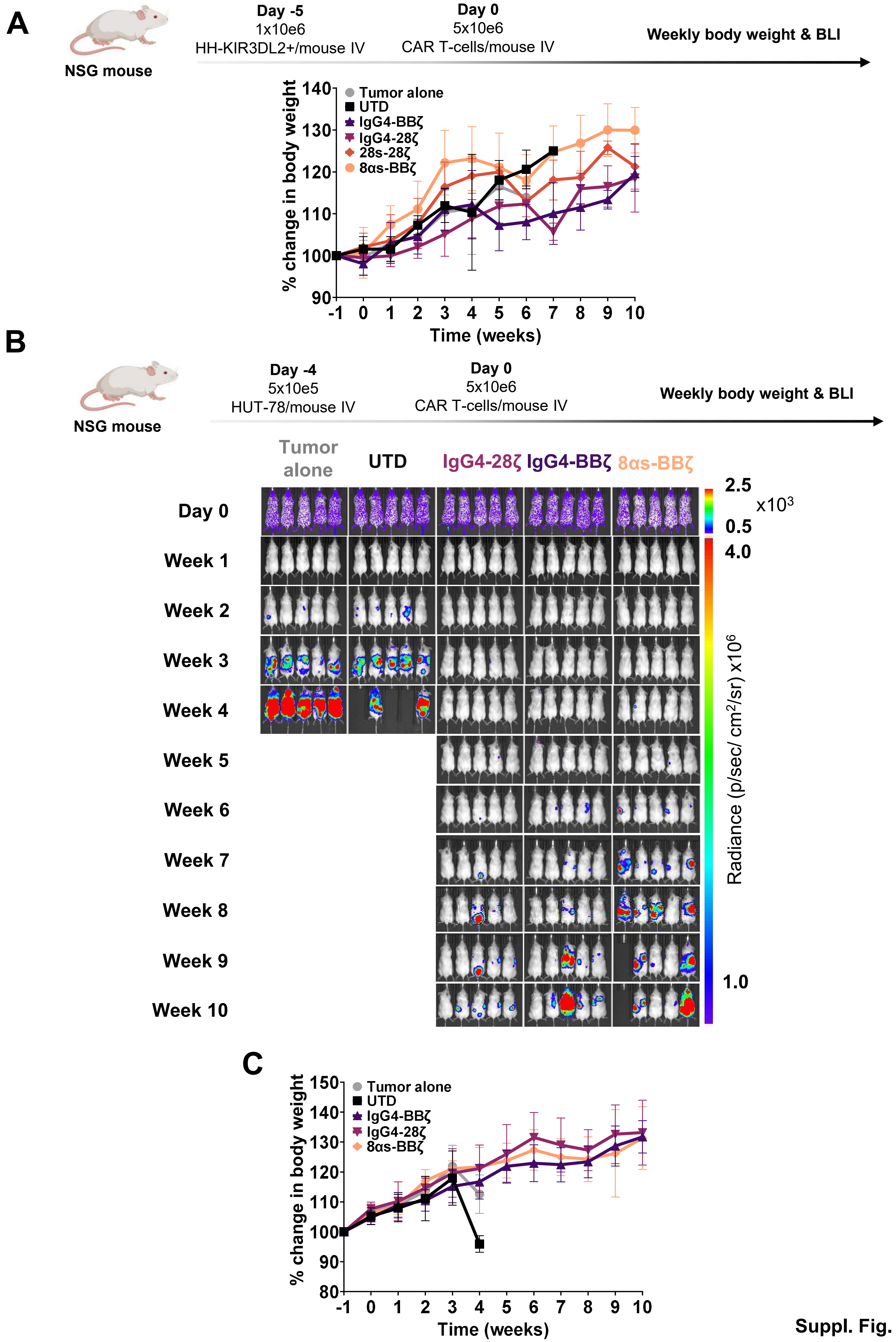
A
NSG mouse
Day -5
1x10e6
HH-KIR3DL2+/mouse IV
Day 0
5x10e6
CAR T-cells/mouse IV
Weekly body weight & BLI
% change in body weight
Time (weeks)
Tumor alone
UTD
IgG4-BBζ
IgG4-28ζ
28s-28ζ
8αs-BBζ
B
NSG mouse
Day -4
5x10e5
HUT-78/mouse IV
Day 0
5x10e6
CAR T-cells/mouse IV
Weekly body weight & BLI
Tumor alone
UTD
IgG4-28ζ
IgG4-BBζ
8αs-BBζ
Day 0
Week 1
Week 2
Week 3
Week 4
Week 5
Week 6
Week 7
Week 8
Week 9
Week 10
2.5
x10³
0.5
4.0
Radiance (p/sec/ cm²/sr) x10⁶
1.0
C
% change in body weight
Time (weeks)
Tumor alone
UTD
IgG4-BBζ
IgG4-28ζ
8αs-BBζ

Suppl. Fig. 4

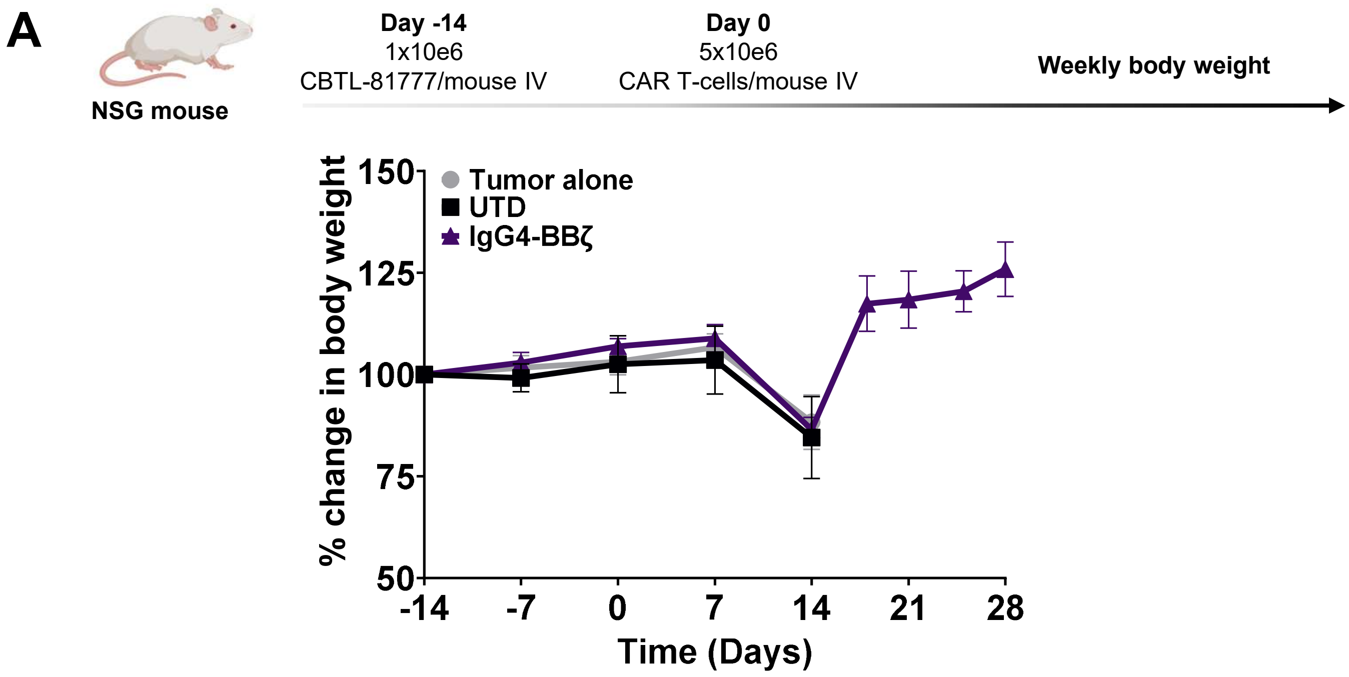
A
NSG mouse
Day -14
1x10e6
CBTL-81777/mouse IV
Day 0
5x10e6
CAR T-cells/mouse IV
Weekly body weight
Tumor alone
UTD
IgG4-BBζ
% change in body weight
150
125
100
75
50
-14
-7
0
7
14
21
28
Time (Days)

Suppl. Fig. 5

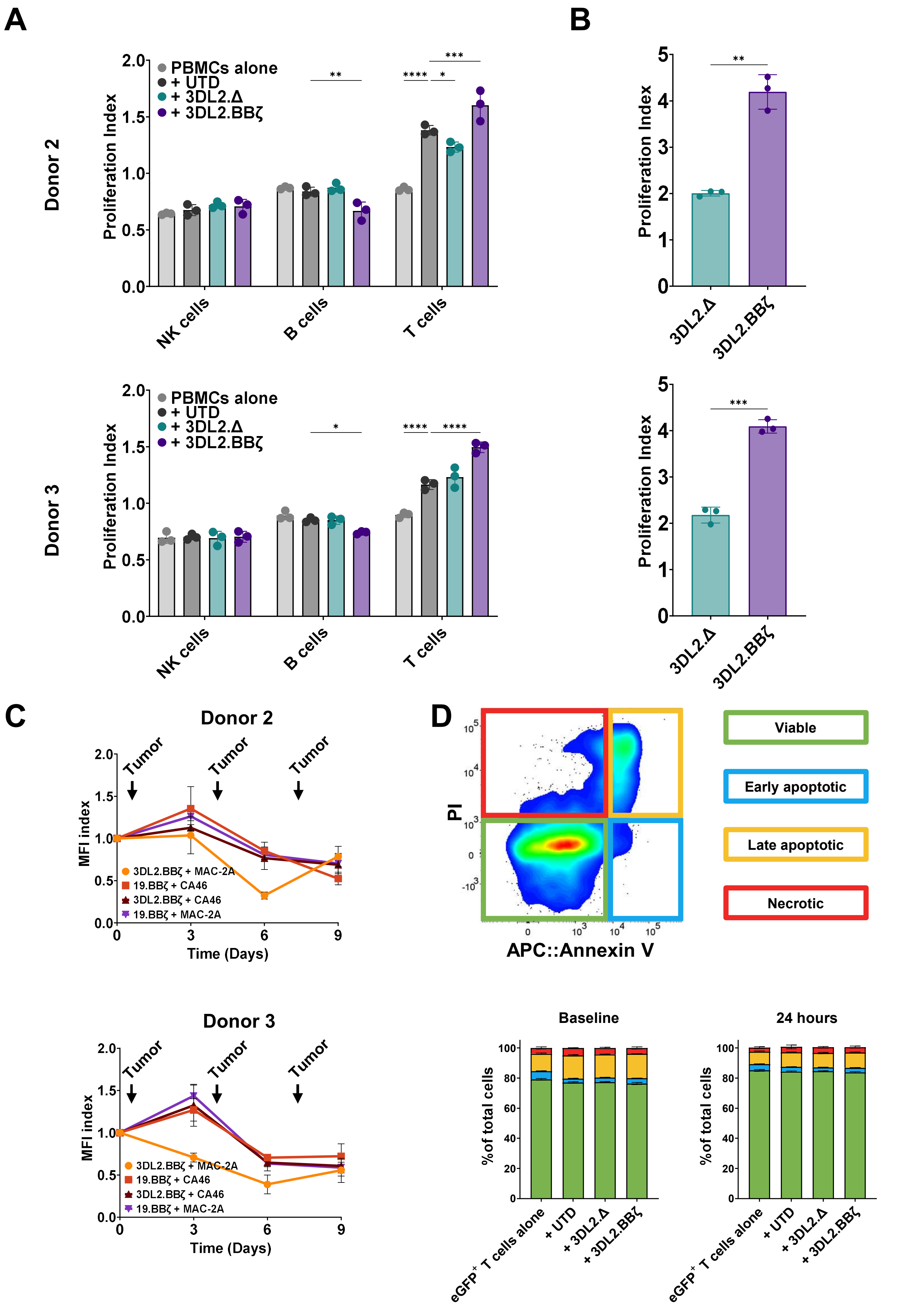
A
B
Donor 2
Donor 3
Proliferation Index
PBMCs alone
+ UTD
+ 3DL2.Δ
+ 3DL2.BBζ
NK cells
B cells
T cells
3DL2.Δ
3DL2.BBζ
C
D
Tumor
MFI index
Time (Days)
3DL2.BBζ + MAC-2A
19.BBζ + CA46
3DL2.BBζ + CA46
19.BBζ + MAC-2A
PI
APC::Annexin V
Viable
Early apoptotic
Late apoptotic
Necrotic
Baseline
24 hours
%of total cells
eGFP+ T cells alone

Suppl. Fig. 6
